# Steering ecological-evolutionary dynamics during artificial selection of microbial communities

**DOI:** 10.1101/264697

**Authors:** Li Xie, Wenying Shou

**Author notes:** Author of correspondence (LX); (WS).

## Abstract

Microbial communities often perform important functions that arise from interactions among member species. Community functions can be improved via artificial selection: Many communities are repeatedly grown, mutations arise, and communities with the highest desired function are chosen to reproduce where each is partitioned into multiple offspring communities for the next cycle. Since selection efficacy is often unimpressive in published experiments and since multiple experimental parameters need to be tuned, we sought to use computer simulations to learn how to design effective selection strategies. We simulated community selection to improve a community function that requires two species and imposes a fitness cost on one of the species. This simplified case allowed us to distill community function down to two fundamental and orthogonal components: a heritable determinant and a nonheritable determinant. We then visualize a “community function landscape” relating community function to these two determinants, and demonstrate that the evolutionary trajectory on the landscape is restricted along a path designated by ecological interactions. This path can prevent the attainment of maximal community function, and trap communities in landscape locations where community function has low heritability. Exploiting these observations, we devise a species spiking approach to shift the path to improve community function heritability and consequently selection efficacy. We show that our approach is applicable to communities with complex and unknown function landscapes.

## Introduction

Multi-species microbial communities often display *community functions* — biochemical activities not achievable by any member species alone. For example, a community of *Desulfovibrio vulgaris* and *Methanococcus maripaludis*, but not either species alone, converts lactate to methane in the absence of sulfate [1]. Community function arises from “interactions” where one community member influences the physiology of other community members. Interactions are typically complex and difficult to characterize, making it challenging to design communities with desired functions [2, 3]. In a different approach, one could mutagenize individual community members, assemble them at various ratios, and screen the resultant communities for high community function. However, this requires community members being culturable, and the number of combinatorial possibilities increases rapidly with the number of species and genotypes. In addition, such assembled communities might be vulnerable to ecological invasion [4].

Alternatively, community function may be improved by top-down artificial selection [5, 6, 7]. During each cycle of artificial community selection, newly-assembled Newborn communities (“Newborns”) grow into Adult communities (“Adults”) for a period of “maturation” time set by the experimentalist. During community maturation, community members can proliferate and mutate. At the end of community maturation, Adults expressing the highest community function are chosen to “reproduce” where each is randomly partitioned into multiple Newborns for the next selection cycle (Figure 1A). Artificial community selection, if successful, can improve useful community functions such as fighting pathogens [8], producing drugs [9], or degrading wastes [10].

**Figure 1:**
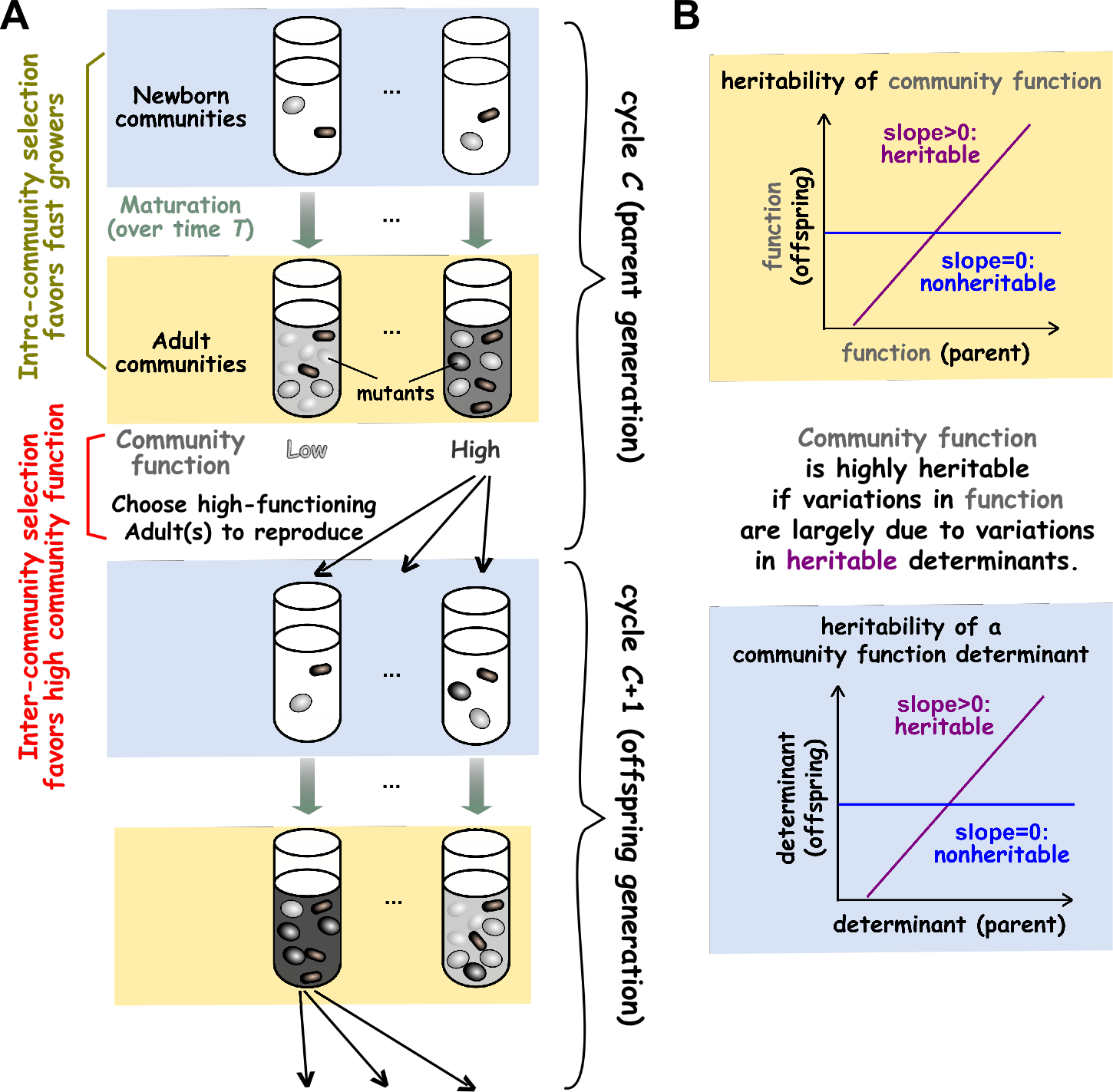
Artificial community selection to improve a community function. **(A)** Artificial community selection. In each cycle, Newborns (blue shading) mature into Adults (yellow shading) during which intracommunity selection (olive) favors fast growers. The highest-functioning Adults are then chosen to reproduce, and this inter-community selection (red) favors high function. Note that both parent (top half) and offspring (bottom half) communities have Newborn and Adult stages. Thus, the four rows from top to bottom represent parent Newborn, parent Adult, offspring Newborn, and offspring Adult, respectively. **(B)** Heritability of community function. Community function is heritable if parent function and average offspring function are positively correlated (top). Community function is highly heritable if variations in function are largely attributable to variations in the heritable determinants (bottom). Heritability can be quantified by the slope of the least squares regression between the parent variable and the average offspring variable. Note that while community function is quantified at the Adult stage, community function determinants can be quantified at the Newborn stage if newly-arising mutations have negligible effects on community function within a cycle.

Theoretical work predicts that artificial selection of communities can succeed, at least under certain conditions [11, 12, 13, 14, 15, 16, 4]. Experimental work on community selection have yielded variable outcomes [17, 18, 19, 20, 21, 22, 23, 24, 25, 26, 27, 28]. In some cases, communities indeed responded to selection, presumably driven by changes in the species composition [22, 23, 24, 25] and/or evolution [17, 18], although some of these experiments are not conclusive due to lack of a “no selection” control. In other cases, selecting for high community function yielded similar outcomes as selecting for low community function or random selection [19, 27, 28, 26], and community function could even decline despite selection [20, 25]. For example, selecting marine microbial communities for enhanced chitin-degradation activity was ineffective unless community maturation time was progressively adjusted to prevent undesirable species from taking over [25].

Artificial community selection is challenging. Unlike artificial selection on individuals, artificial community selection is influenced by inter-species ecological interactions, and species coexistence must be ensured. Artificial community selection is particularly challenging when community function incurs a fitness cost to one or more community members. This cost can take diverse forms. For example, engineered microbes must divert cellular resources to make a product desired by experimentalists, which can slow down cell growth. As another example, fast-growing species in a community must evolve to grow slower to not outcompete their partners. When community function is costly, cheaters who do not contribute to community function will grow faster than cooperators who contribute. Thus, cheaters are favored by intra-community selection during community maturation (Figure 1, olive). However, cheaters are disfavored at the end of a selection cycle by inter-community selection that only allows communities with high functions to reproduce (Figure 1, red). Thus, to improve a costly community function, inter-community selection must overwhelm intra-community selection (Figure 1). Indeed, simulations and experiments have shown that artificial community selection could fail unless a selection scheme takes into consideration a variety of factors, including promoting species coexistence, suppressing cheaters, choosing an appropriate number of Adult communities to reproduce, and introducing heritable variations while reducing nonheritable variations in community function [6, 15, 25, 26, 28, 4].

Successful community selection requires three elements: variations in community function (enabled by mutations or species migrations for example), preferential survival of high-functioning communities (enabled by inter-community selection), and heritability of community function [29]. Community function is heritable if parent function is positively correlated with the average offspring function (Figure 1B, top). Community function has high heritability if most variations in community function can be attributed to variations in heritable determinants (Figure 1B, bottom). Community function determinants (“determinants”) include, for example, the species and genotype compositions of a community. These determinants can be defined at the Newborn stage if any newly-arising genotypes (e.g. cheaters) cannot rise to a sufficient level to affect community function within that cycle. Some community function determinants are heritable to some extent (e.g. genotype compositions), while others are not heritable (e.g. stochastic fluctuations in species compositions due to, for example, pipetting) [15].

In this article, we focus on the heritability of community function and how it is affected by ecological species interactions. Using computer simulations, we investigate artificial selection of two-species commensal communities. Computer simulations allow us to compare multiple selection strategies, and to learn insights that will guide future experiments. With a two-species community, we can boil community function down to its most important, and orthogonal, components: a heritable determinant and a nonheritable determinant. This in turn enables us to visualize a “community function landscape” relating community function to its heritable and nonheritable determinants, similar to a phenotype landscape relating an individual’s phenotype to its genetic and environmental determinants [30, 31]. The local geometry of the landscape can then be used to visualize the heritability of community function, an idea similar to phenotype landscape indicative of heritability of individual traits. Within this conceptual framework, we find that the very effort of promoting species coexistence can constraint selection efficacy. Specifically, we promoted species coexistence by using commensalism [32], which is common in microbial communities [33, 34, 35, 36, 37, 38]. However, the ensuing steady state species composition, visualized by the species composition “attractor”, means that evolving communities can only sample a small region of the landscape. This can prevent the attainment of maximal community function, and trap communities in landscape locations where community function has low heritability. Inspired by these observations, we devise selection strategies that improve heritability and selection outcomes, even when the community function landscape is high-dimensional and unknown.

## Results

### A commensal Helper-Manufacturer community displays a species composition “attractor”

In our previous work [15], we simulated artificial selection on a two-species Helper-Manufacturer community (“H-M community”, Figure 2A). In this community, Helper (H) digests agricultural waste and consumes Resource, grows biomass and, at no cost to itself, releases a metabolic Byproduct essential for Manufacturer (M). As M consumes Byproduct and Resource, its cellular resource is partitioned so that a fraction *f_P_* (0 ≤ *f_p_* ≤ 1) is used to synthesize a high-value product (Product *P*), while the rest (1 − *f_P_*) is used for its own biomass accumulation.

**Figure 2:**
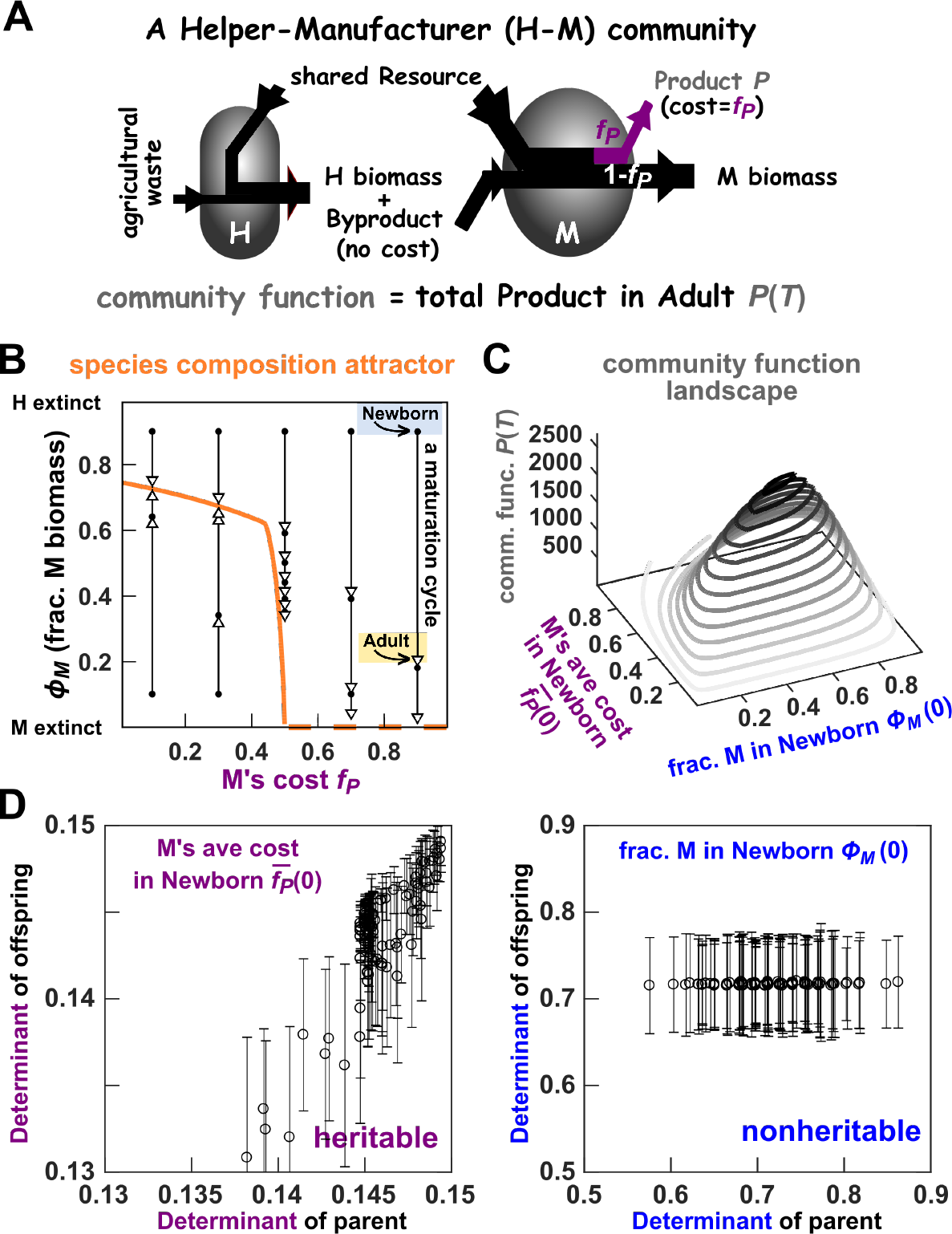
Ecological dynamics and community function landscape of a two-species commensal community. **(A)** The Helper-Manufacturer commensal community. **(B)** Species composition attractor. Each arrow describes the change in *ϕ_M_*, the fraction of M’s biomass in a community, from Newborn (solid dot) to Adult (open triangle) over one maturation cycle. All arrows are “attracted” to the species composition attractor (orange curve). When M’s cost *f_P_* is high, M goes extinct (broken orange line), and at lower *f_P_* values, M coexists with H (solid orange line). **(C)** Community function landscape. Community function *P*(*T*) is plotted in 3D as a function of Newborn community’s fraction of M biomass (*ϕ_M_* (0)) and average cost among M 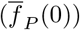. To ensure that community function only has these two determinants, we made the following simplifications: i) Newborn’s total biomass and the initial abiotic environment are both fixed; ii) stochastic death rate is relatively low to ensure largely deterministic population dynamics; and iii) mutation rate (0.002/cell/generation) and community maturation time *T* (~6 population doublings) are relatively small so that newly-arising genotypes remain rare within a cycle and not impact community function (Figure S7 of [15]). Indeed, community functions observed in simulations are well predicted by those calculated from the two determinants *ϕ_M_* (0) and 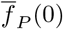 (Figure S1). Species composition attractor and community function landscape were calculated using Eqs. 1–5 with the Newborn total biomass=100 and maturation time *T* = 17. **(D)** Newborn average cost 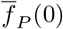 is largely heritable (positive slope), but Newborn’s fraction of M biomass *ϕ_M_*(0) is not heritable (zero slope). Each open circle is obtained by averaging over ~60 offspring Newborns descending from the same parent community. Error bars indicate one standard deviation. Note that for 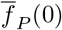, linear regression did not pass through the origin. Offspring costs tend to be less than parent costs because during community maturation, the average cost declines as cheaters take over.

Since M relies on H’s Byproduct, H can either drive M extinct or coexist with M [15]. When M’s cost *f_P_* is large, M always grows slower than H. Thus, *ϕ_M_*, the fraction of M biomass in a community, declines within a cycle and over cycles until M goes extinct (Figure 2B, dashed orange line). When M’s cost *f_p_* is moderate or small, and when we choose growth parameters of the two species properly (Table 1), H and M can coexist at a steady state ratio (Figure 2B, solid orange line). Note that communities with stable coexistence of species have been engineered in the lab (e.g. [39, 40]). Steady state fraction of M biomass in a community at various cost *f_p_* form a species composition “attractor”: at a given cost, species composition away from the attractor is pulled toward the attractor as the community matures (arrows pointing toward the orange curve in Figure 2B).

**Table 1:** Parameters for phenotypes of H and M used in the simulations. Simulations in Figure 7 start with H and M whose phenotypes are listed in the “Ancestral” column. These phenotypes are not allowed to change beyond values listed in the “Bounds” column. In most simulations where only *f_p_* can be modified by mutations, except for *f_P_*, parameters in the Bounds column are used. For maximal growth rates, * represents evolutionary upper bound. For *K_SpeciesMetabolite_*, * represents evolutionary lower bound, which corresponds to evolutionary upper bound for Species’s affinity for Metabolite (1/*K_SpeciesMetabolite_*). For parameter justifications, see [15].

|  | Definition | Ancestral | Bounds |
| --- | --- | --- | --- |
| $R(0)$ | initial amount of Resource in Newborn | 1 | |
| $f_P$ | fraction of M growth diverted to producing P | 0.10 | 1 |
| $K_{MR}$ | fold of $R(0)$ at which $g_{Mmax}/2$ is achieved in excess B | 1 | $1/3^*$ |
| $K_{MB}$ | amount of Byproduct at which $g_{Mmax}/2$ is achieved in excess R | $\frac{5}{3} \times 10^2$ | $\frac{1}{3} \times 10^{2*}$ |
| $K_{HR}$ | fold of $R(0)$ at which $g_{Hmax}/2$ is achieved | 1 | $1/5^*$ |
| $g_{Mmax}$ | maximal biomass growth rate of M | 0.58/unit time | 0.7/unit time* |
| $g_{Hmax}$ | maximal biomass growth rate of H | 0.25/unit time | 0.3/unit time* |
| $\delta_M$ | death rate of M | $3.5 \times 10^{-3}$/unit time | |
| $\delta_H$ | death rate of H | $1.5 \times 10^{-3}$/unit time | |
| $c_{RM}$ | fraction of $R(0)$ consumed per M biomass grown | $10^{-4}$ | |
| $c_{RH}$ | fraction of $R(0)$ consumed per H biomass grown | $10^{-4}$ | |
| $c_{BM}$ | amount of Product consumed per M biomass grown | $\frac{1}{3}$ | |
| $P_{mut}$ | mutation probability per cell division for each mutable phenotype | $2 \times 10^{-3}$ | |

### Visualizing community function landscape in relation to heritable and nonheritable determinants

H-M community function is defined as the total amount of Product accumulated in an Adult community, denoted by *P*(*T*) with *T* being the community maturation time. Community function requires both species, since H supports M while M makes Product. Community function is not costly to H, but costly to M (cost = *f_P_*).

What determines community function? If we allow only M’s cost *f_p_* to mutate and make some additional simplifications (details in Figure 2C legend), community function *P*(*T*) can be adequately predicted from two determinants: the fraction of M biomass in the Newborn (*ϕ_M_*(0)), and the average cost paid by M in the Newborn 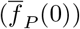. We can now visualize “community function landscape” in a 3D plot of community function *P*(*T*) as a function of its two determinants, similar to a topographic map (Figure 2C). The single peak corresponds to a global maximal community function achieved at an intermediate species composition and an intermediate M cost [15]. Note that when M pays a low cost, community function is low because little product is made, but when M pays a very high cost, community function is also low because M cannot proliferate substantially (Figure 2B). Similarly, community function peaks at an intermediate value of Newborn species composition (*ϕ_M_*(0)), since community function requires both M and H. Thus, we have simplified our system so that community function has only two determinants.

Of the two determinants, M’s average cost 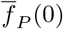 is heritable, while Newborn species composition *ϕ_M_*(0) is non-heritable (Figure 2D). For 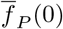, if a parent Newborn is dominated by cheaters (low 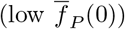) or cooperators (high 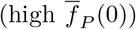), then so will its offspring Newborns (Figure 2D left). However, offspring costs are generally less than the parent cost, since cooperator frequency declines during community maturation due to cheater takeover (Figure 2D left). For *ϕ_M_*(0), when we generate Newborns in the parent cycle, species composition can fluctuate stochastically (e.g. a species ratio of 50:50 can become 40:60 by chance due to pipetting a small number of cells). However, these stochastic fluctuations are dampened as species compositions gravitate toward the attractor during community maturation (Fig 2B, solid orange). These Adults end up sharing similar species compositions, and so will their offspring Newborns. In essence, variations in Newborn species composition are not transmitted across cycles, and therefore are not heritable (Figure 2D, right).

In effective selection, variations in nonheritable determinants should be minimized, since any elevation in community function due to nonheritable determinants will not transmit to the next generation. Thus, we can quantify selection efficacy as the progress in the heritable determinant 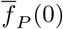.

### Constrained by attractor, community selection fails to reach maximal function

We now consider the case where ancestral M pays a cost smaller than what is optimal for maximal community function. While inter-community selection will favor cooperator M paying a higher cost, intra-community selection will always favor cheater M paying a lower cost (Figure 1A). If we randomly select Adults to reproduce, or allow each Adult to reproduce one offspring, then community function rapidly declines to zero as cheaters take over [15]. If we apply community selection, can we improve community function to the maximal value?

We find that although community selection can improve community function, community function fails to reach the maximum ([15]; Figure 3A), even after variations in the nonheritable determinant *ϕ_M_*(0) have been minimized. We minimized variations in nonheritable determinant by reproducing Adults via “cell sorting”, as if by a flow sorter capable of measuring the biomass of individual H and M cells so that all Newborns had nearly identical species composition. With this experimentally-challenging community reproduction method, community selection successfully improved community function over ~200 selection cycles (Figure 3A). However, community function *P*(*T*) never reached the theoretical maximum (Figure 3A dashed line). The average cost paid by M 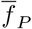 stayed above the optimal 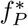 (Figure 3B), which is surprising because high cost would be disfavored by both intra-community selection (which favors fast growth and low cost) and inter-community selection (which favors 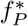).

**Figure 3:**
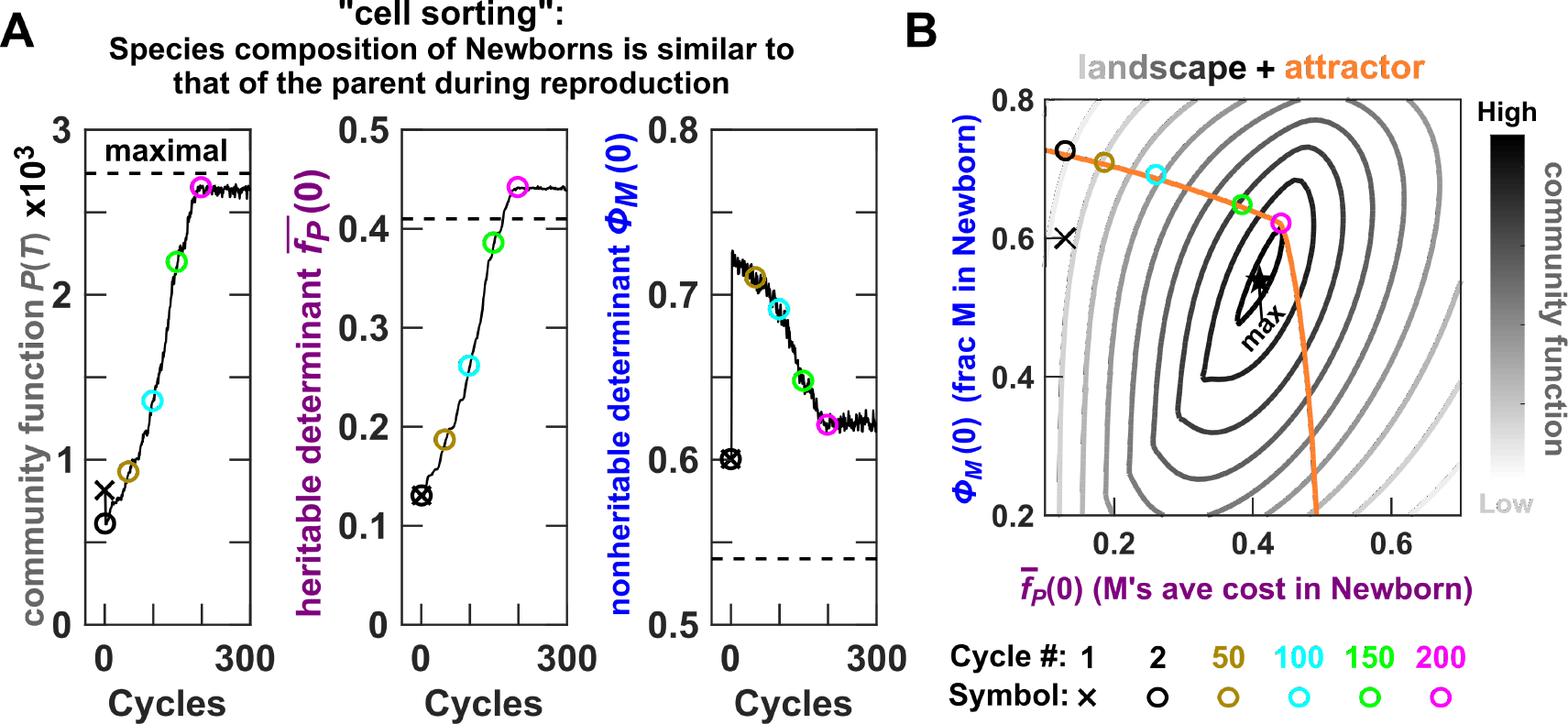
Species composition attractor prevents the attainment of maximal community function. **(A)** The evolutionary dynamics of the H-M community under a “cell sorting” community selection scheme (Methods; Fig 3D of Ref. [15]). A selection cycle began with a total of 100 Newborn communities, and two communities with the highest amount of Product were chosen to reproduce where each was randomly split into 50 Newborns. Chosen Adults were reproduced through cell sorting, and thus each Newborn had a total biomass very close to the target value of 100 and species ratio very close to that of the parent Adult. If species ratio was allowed to fluctuate stochastically as in pipetting, community function failed to improve (Figure S12 leftmost panel in Ref. [15]). Community function determinants 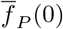 or *ϕ_M_* (0) were obtained from the Newborn stage of chosen Adults, and then averaged over chosen communities. The black dashed lines mark the theoretical maximum of *P*(*T*) and corresponding optimal 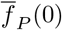 and *ϕ_M_* (0). **(B)** Species composition attractor constrains community function away from the maximal. Evolutionary dynamics of community function determinants are plotted on top of community function landscape overlaid with the species composition attractor. The first cycle is marked with a cross and representative subsequent cycles are marked with circles.

To understand this sub-optimal selection outcome, we overlaid the community function landscape with the species composition attractor (steady state species compositions at various cost values). We then plotted the evolutionary dynamics of community function determinants (Figure 3B circles). The evolutionary dynamics were clearly confined to the species composition attractor (circles on top of the orange line), and that the attractor did not pass through the maximal community function (orange line not crossing the black star). This explains the sub-optimal selection outcome: Like a hiker who is restricted to a trail that does not traverse the mountain top, community function can only climb to the highest value along the attractor (magenta circle), which is lower than the global maximum (black star). Consequently, the corresponding 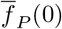 and *ϕ_M_*(0) are higher than their respective optima. In other words, selection outcome is constrained along the species composition attractor by ecological forces.

### The local geometry of community function landscape predicts the efficacy of inter-community selection

The local geometry of community function landscape predicts how much inter-community selection can advance heritable determinant toward the theoretical optimum. Specifically, in the region of the landscape where community function contour lines are perpendicular to the axis of the heritable determinant (Figure 4A), variation in community function is fully attributed to variation in the heritable determinant. Thus in such regions, community function is heritable, and inter-community selection can make maximal progress in the heritable determinant. In contrast, when community function contour lines are parallel to the axis of the heritable determinant (Figure 4B), no variation in community function can be attributed to variation in the heritable determinant. Thus, community function is nonheritable, and inter-community selection makes no progress in the heritable determinant. An intermediate case is shown in Figure 4C.

**Figure 4:**
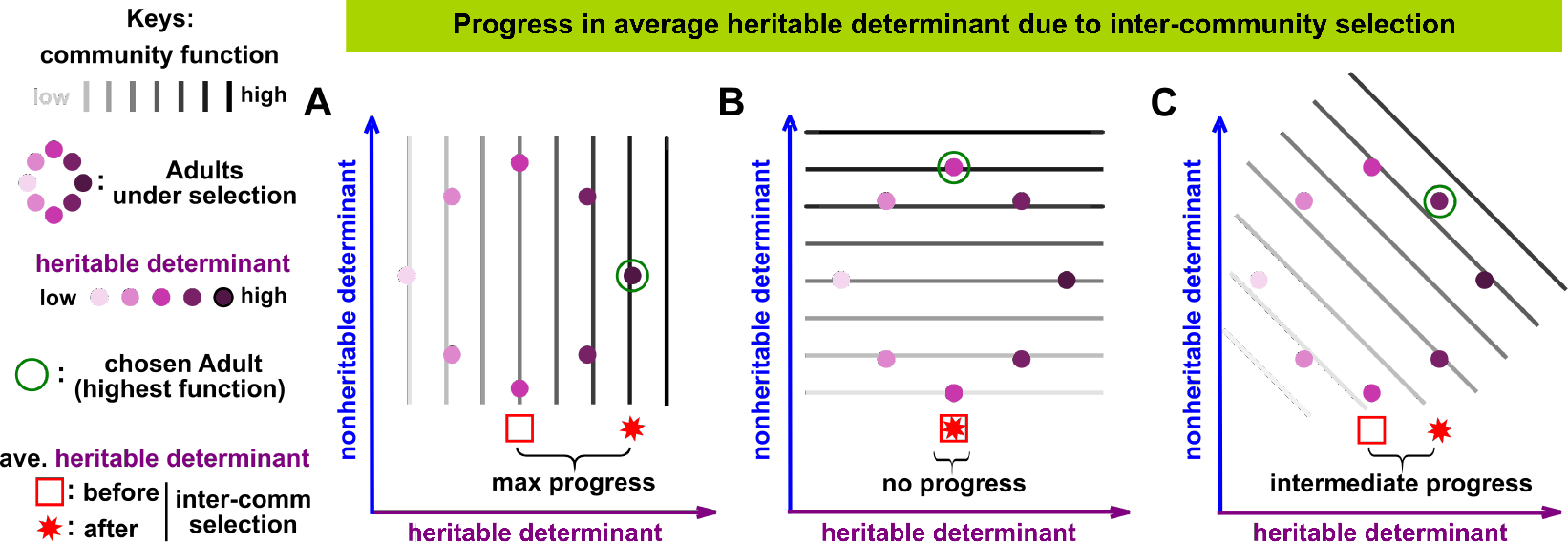
The efficacy of inter-community selection depends on the local geometry of community function landscape. Community function landscape is plotted in contours where darker shade indicates higher function. Eight Adults are under selection (dots with light to dark purple shades reflecting low to high heritable determinant). The community with the highest function is chosen for reproduction (green open circle). Community selection is the most or the least effective when community function contour lines are perpendicular **(A)** or parallel **(B)** to the axis of heritable determinant, respectively, and intermediate otherwise **(C)**. Average heritable determinant before and after inter-community selection are marked by a red square and a red star, respectively. Thus, progress made by inter-community selection can be measured as the distance from red square to red star.

### Species spiking can improve selection outcome by shifting the location of attractor

We can now examine the efficacy of H-M community selection within community function landscape (Figure 5Bi and ii). We allowed Newborn species composition to stochastically fluctuate, as if pipetting samples from an Adult to seed Newborns while keeping the total Newborn turbidity fixed. This experimentally-facile community reproduction method resulted in a “cloud” of Newborns around the species composition attractor (Figure 5Bii), and their predicted community functions at Adulthood could be read out from the landscape based on the values (shade) of the contours. Although landscape contours near the attractor were not straight lines, they were largely parallel to the heritable determinant axis (similar to Figure 4B). Thus, variations in community function were largely attributed to variations in the nonheritable determinant. Indeed community function had low heritability (Figure 5Cii), and inter-community selection made only small progress in the heritable determinant (short red arrow in Figure 5Bii; statistics in the red box plot of Figure 5Biv). This small progress was just enough to counter the decline due to intra-community selection (olive box in Figure 5iv), resulting in a net of zero progress (black box in Figure 5iv). Consequently, despite 1000 selection cycles, average community function and average heritable determinant among chosen communities barely improved (Figure 6A).

**Figure 5:**
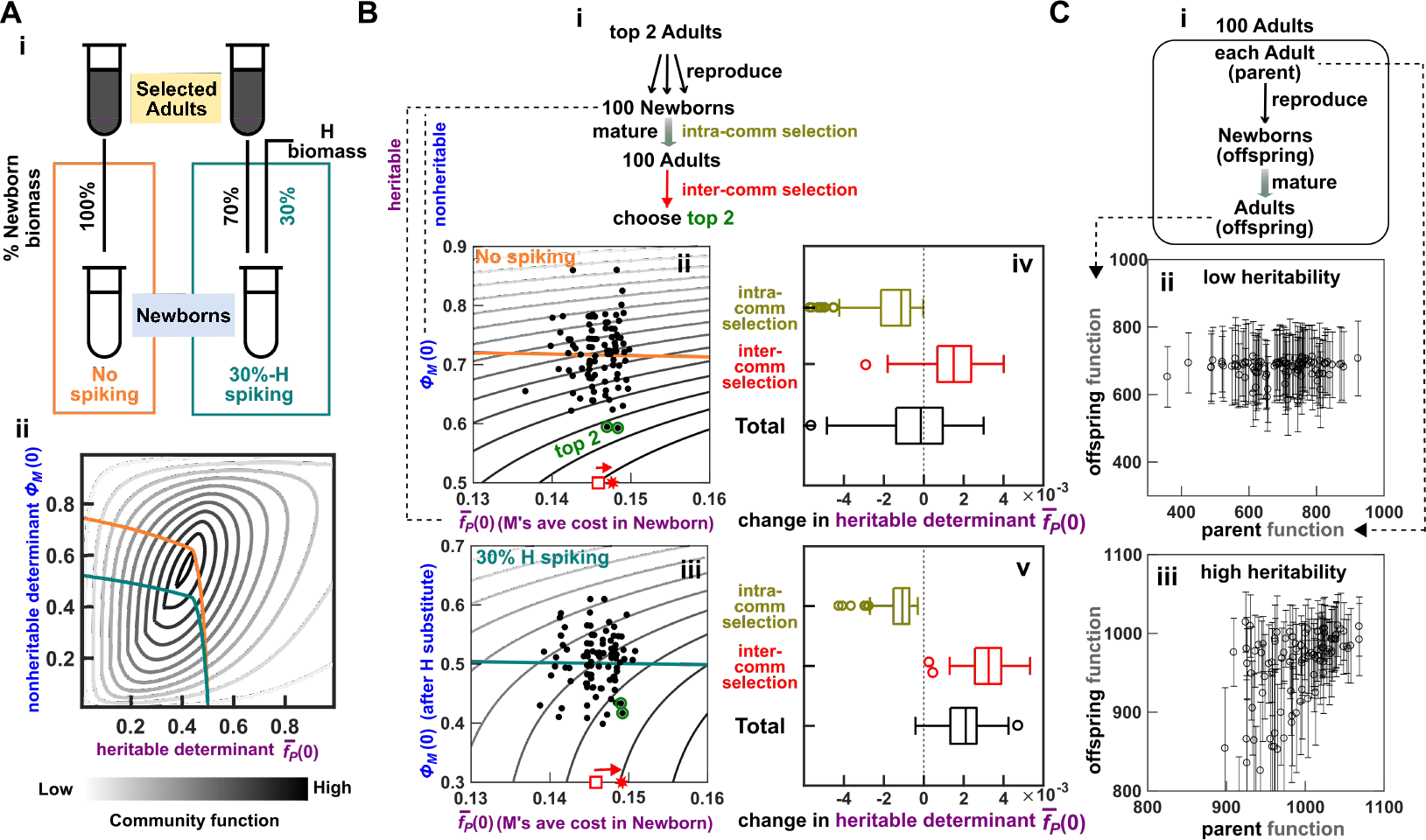
Shifting the species composition attractor to a favorable region in landscape increases community function heritability and selection efficacy. **(A)** Species spiking shifts attractor. Species composition attractor (orange curve in ii) is shifted when 30% of the total biomass in a Newborn is always replaced by the nonevolving H (“30%-H spiking”; teal curve in ii). **(B)** Community selection outcome can be significantly improved by species spiking. **i**: Scheme of community selection over one cycle. **ii, iii**: Visualization of inter-community selection on the landscape under the no-spiking strategy and the 30%-H spiking strategy, respectively. Because communities are constrained by the attractor, we only need to focus on the geometry of the landscape near the attractor. The top 2 communities chosen to reproduce are highlighted with green circles. The average heritable determinant 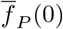 over all 100 communities and over the top 2 are indicated by red open square and red star, respectively. The length of the red arrow indicates the progress in heritable determinant 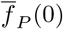 due to inter-community selection. **iv, v**: Progress in the heritable determinant 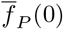. Selection cycle was repeated 100 times to obtain box plots (center: median; bottom and top edges: the 25th and 75th percentiles, respectively; whiskers: data range excluding outliers; open circles: outliers). Although inter-community selection (red) improves heritable determinant 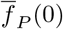, intra-community selection (olive) reduces it. The sum of these two effects is the net progress (black). **(C)** Species spiking can improve the heritability of community function. **i**: Heritability is quantified by comparing 100 pairs of parent functions and their average offspring functions and estimating the slope of the least squares linear regression. **ii**: Under the no-spiking strategy, community function heritability is low. **iii**: Under the 30%-H spiking strategy, community function heritability is high. Error bars indicate one standard deviation. Empirical determination of this relationship between offspring and parent function can be used in lieu of landscape visualization to guide spiking strategy.

**Figure 6:**
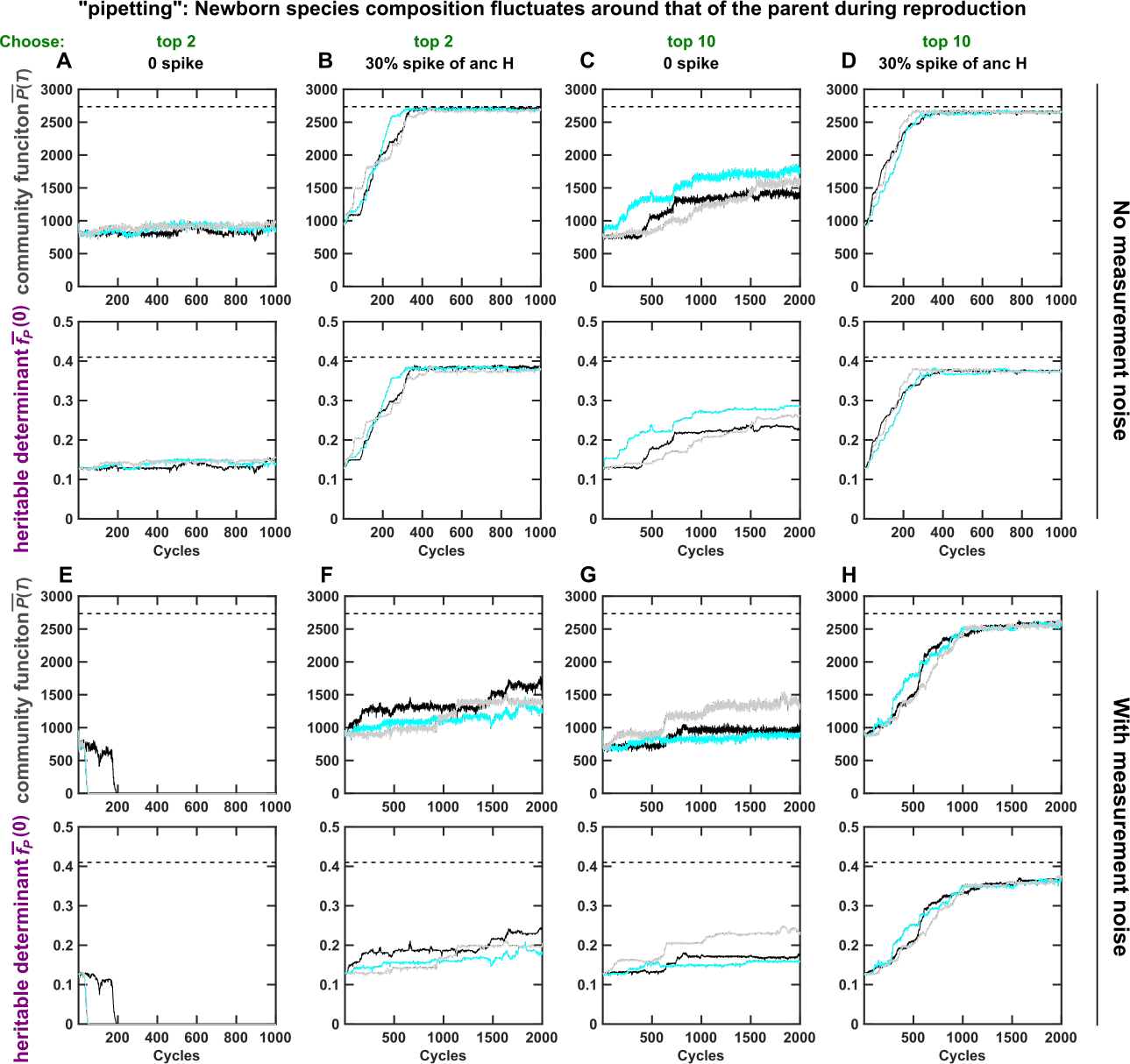
Evolution dynamics of community function *P*(*T*) and its heritable determinant 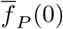 averaged over the chosen communities. Species composition but not total biomass of Newborns was allowed to stochastically fluctuate during community reproduction, as if pipetting an inoculum of a fixed turbidity from an Adult to seed its Newborns. In all cases, community selection is more effective under the 30%-H spiking strategy (where 30% of Newborn biomass was replaced by the non-evolving H) compared to the no-spiking strategy. In “top-2” strategy, top 2 communities each reproduced 50 Newborns. In “top-10” strategy, top 10 communities each reproduced 10 Newborns. In “with measurement noise”, measured community function was the sum of the real community function and a normal random variable with a mean of 0 and a standard deviation of 100. Dashed lines correspond to values optimal for community function.

Since the contour of landscape around the attractor was not conducive to effective selection, we improved selection by shifting the attractor to a more favorable location. Specifically, when we replaced 30% of total Newborn biomass with ancestral non-evolving H (“30%-H spiking”, teal box in Figure 5A i), the attractor was forced into a region where community function contour lines were largely perpendicular to the axis of heritable determinant (Figure 5Aii). Community function became more heritable (Figure 5C iii). Inter-community selection made a larger improvement in the heritable determinant (longer red arrow in Figure 5Biii compared to Bii; red box in Figure 5Bv), and total progress was positive (black box in Figure 5Bv). Consequently, community selection was effective (Figure 6B).

The 30%-H spiking strategy also improved selection efficacy when top 10, instead of top 2 communities were selected for reproduction (Figure 6 C vs. D), and when selection suffered interference from measurement noise in community function (Figure 6 E vs. F; G vs. H). The spiking strategy worked even if Newborn species composition and total biomass were both allowed to fluctuate as if the chosen Adults were reproduced and spiked through pipetting (Figure S3). Although quantitative details differ, H spiking at a wide range of percentages improved selection outcomes (Figure S3).

### Dynamic species spiking based on maximizing community function heritability improves selection outcome on a high-dimensional landscape

So far, we have chosen a simple scenario with two determinants so that we can visualize community function landscape. However, landscapes of most community functions are high dimensional and unknown. It is thus infeasible to devise a spiking strategy based on landscape visualization. However, landscape geometry is reflected in the heritability of community function (Figure 5) which can also be estimated from experimental measurements (Figure 5C). Thus, we can try different spiking strategies, and choose the strategy corresponding to the highest community function heritability. Since communities move in the landscape as they evolve, periodic heritability evaluation is needed to dynamically update spiking strategy.

To demonstrate dynamic spiking strategy, let’s consider a more complex scenario with the H-M community. If we allow growth parameters of H and M to also evolve, then community function will have 6 heritable determinants all defined at the Newborn stage: M’s average cost, the average maximal growth rate of H and of M, the average affinities of M to Resource and to Byproduct, and the average affinity of H to Resource. If we reproduce the Adult via “pipetting”, then Newborn’s total biomass and species composition will fluctuate stochastically, adding two nonheritable determinants. If we additionally consider community function measurement noise (a normal random variable with mean 0 and standard deviation comparable to ancestral community function), we will have yet another nonheritable determinant. Overall, community function will have 9 determinants, 6 heritable and 3 nonheritable.

We performed community selection in the above complex scenario (schematic in Figure S4). We started with the no-spiking strategy, and always chose top 10 Adults where each reproduced 10 Newborns (since choosing top 10 worked better than choosing top 2; Figure 6; Ref. [15]). If a fraction of Newborn biomass is to be replaced with H (or M) biomass, the spiking mix consisted of equal parts of 5 evolved H (or M) clones randomly isolated from the previous cycle of the same lineage (Figure S4A). During reproduction, portions of a chosen Adult and the spiking mix were “pipetted” to initiate Newborns so that both the total biomass and the species composition fluctuated stochastically in Newborns.

Every 100 cycles, we evaluated spiking strategy, and updated the strategy when appropriate. Specifically, we quantified community function heritability for 5 candidate spiking strategies (no spiking, 30%-H spiking, 60%-H spiking, 30%-M spiking and 60%-M spiking) by regressing parent function with median offspring function (similar to Figure 5C). The current spiking strategy was then updated if an alternative strategy conferred significantly higher community function heritability (Figure S4B, Methods).

With the no-spiking strategy, community selection moderately improved community function and heritable determinants (Figure 7A and B; Figure S5A). Selection outcome was significantly improved when we dynamically adjusted the spiking strategy according to community function heritability (Figure 7C and D; Figure S5B). In contrast, adopting the spiking strategy with the lowest heritability led to much worse selection outcome (Figure S5C), while randomly choosing spiking strategy led to variable results. It is also noteworthy that during dynamic spiking, the adopted spiking strategy was not static (Figure 7E). A static 60%-H or 30%-H spiking strategy offered negligible improvement over no spiking (Figure S6). Therefore, it is important to evaluate heritability periodically and update accordingly.

**Figure 7:**
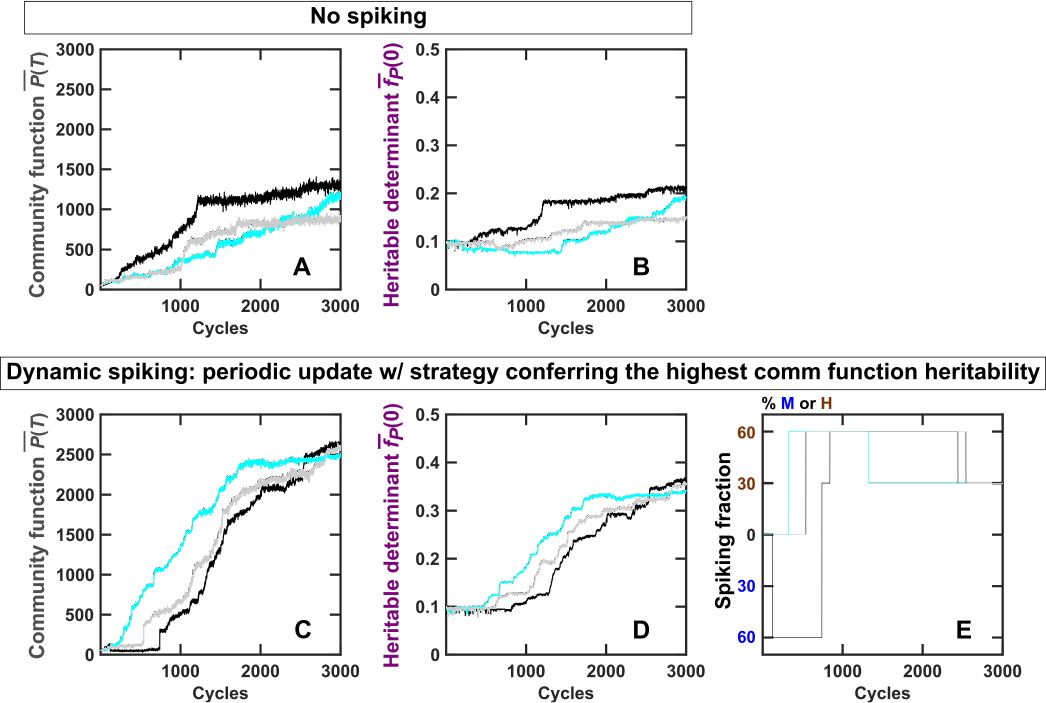
Dynamic species spiking improves selection efficacy. Plots show dynamics of the average community function 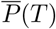 and the average heritable determinant 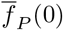 of chosen communities under the no-spiking strategy **(A, B)** and under the dynamic spiking strategy **(C, D)**. During dynamic spiking, 5 candidate spiking strategies (no spiking, 30%-H spiking, 60%-H spiking, 30%-M spiking and 60%-M spiking) were periodically evaluated for community function heritability. The current strategy was replaced by a new strategy if the new strategy conferred significantly higher heritability. Spiking mix consisted of 5 evolved H clones or 5 evolved M clones randomly isolated from the same community lineage in the previous cycle. The corresponding spiking percentages during dynamic spiking are displayed in **(E)**. Black, cyan and gray curves represent 3 independent simulation replicas. Due to the high dimensional landscape, we can no longer determine the theoretical maximal community function.

When we varied parameters of dynamic spiking, selection efficacy did not vary significantly. For example, when the number of colonies used to make the spiking mix was changed from 5 to 1, 2, or 10 clones, qualitatively similar results were obtained (Figures S7, S8 and S9). We also repeated the simulations with 3 candidate spiking strategies: no spiking, 30%-M spiking and 30%-H spiking, and obtained qualitatively similar results (Figure S10). In conclusion, dynamic species spiking based on heritability of community function can improve the efficacy of selection for communities with high-dimensional unknown landscape.

## Discussion

Efficient artificial community selection relies on optimizing inter-community variation, selection strength, and community function heritability. For example, selection outcome significantly improved when we minimized nonheritable variations by fixing Newborn total biomass and species composition instead of allowing them to stochastically fluctuate ([15]; compare Figure 3A with Figure S3A). In Ref. [4], the authors improved selection efficiency by increasing inter-community variation through species migration and population bottleneck. Intriguingly, community trait evolved to be stable even though stability was not directly selected for. This is consistent with the notion that only heritable community trait can respond to selection.

A challenging aspect of optimizing community selection is that variation, selection strength, and heritability are interconnected. For example, strong selection can be counterproductive because it can reduce inter-community variation (compare “top 2” with “top 10” in Figure 6) [15]. During community reproduction, drastically diluting the parent Adult (strong bottleneck) will increase inter-community variation, but will also create large stochastic variations which reduce heritability. Similarly, mixing Adults before reproduction (“migrant pool”) increases variation but at the cost of heritability. Indeed, reproducing H-M communities via the migrant pool method causes poor selection outcome due to the spread of cheaters (Figure S2).

Our current work showcases the difficulty in achieving community function heritability. On the one hand, beneficial species interactions, such as commensalisms and mutualisms, can enhances heritability of community function by ensuring species coexistence [32, 41]. However, the resultant species composition attractor means that without interventions, communities may only sample a small region in the community function landscape. This can constrain both the dynamics (Figure 5) and the long-term outcome of selection (Figure 3B): If the species composition attractor does not pass through the maximal community function, selection outcomes will be suboptimal (Figure 3B). Furthermore, if the species composition attractor sits in a region where community function heritability is low, then selection efficacy will be low.

These concepts have prompted us to devise species spiking strategies that might force the attractor to a more favorable region with higher community function heritability. When we can visualize the landscape, we can design spiking strategy to force the attractor to a favorable region (i.e., where contour lines of community function are perpendicular to the axis of heritable determinant, Figures 4 and 5). When we can not visualize the landscape, we can choose spiking strategy based on community function heritability (Figure 7), a quantity that can be estimated in experiments (Figure 5C). Spiking strategy needs to be dynamically adjusted since evolving communities move in the landscape (Figure S6).

Species spiking has limitations and caveats. First, member species need to be culturable so that we can isolate clones to form spiking mixes. However, flow sorting member species could potentially bypass this problem. Second, evaluating community function heritability is resource-intensive. If a community has multiple species, many spiking strategies may need to be compared. Thus, an important future direction is to understand how to optimize dynamic spiking strategy and to reduce workload. Third, extreme spiking percentages can induce spurious heritability. For example, when we considered only 3 candidate strategies (no spiking, 30%-H spiking, and 30%-M spiking), selection efficacy can be higher than when we also included the 60%-H spiking and 60%-M spiking strategies (Figure S10). This is because 60%-H spiking was so extreme that species composition did not return to the attractor within one cycle (Figure S11). Consequently, stochastic fluctuations in species composition became partially heritable. Positive correlation between nonheritable *ϕ_M_* (0) of the parent and offspring communities thus contributes toward community function heritability, misleading the choice of spiking strategy.

Although here we have considered selection of H-M communities, the ideas of community function landscape, species composition attractor, and heritability are general. Indeed, experiments show that low heritability of community function could limit selection efficacy [26, 28, 27]. We therefore propose that it is possible to devise appropriate spiking strategies for other types of communities, particularly those with a species composition attractor (i.e. stable species composition). Future directions would include testing spiking strategies on diverse communities, both with simulations and in lab experiments.

Over the last century, a rich body of theory has been developed to understand evolution of quantitative traits in individual organisms (e.g. [42]). Two of the key concepts we use here are landscape and heritability, which are fundamental for understanding evolution of individual traits. For example, phenotype landscape as a function of genetic and environmental determinants was used to illustrate the evolution of developmental interactions ([30] and [31]). The local geometry (gradient) of landscape determines how sensitive the phenotype is to underlying variations, and thus to selection based on the phenotype. Heritability is pivotal in evolution biology, particularly in breeding. Multiple statistical methods have been developed to estimate heritability and facilitate designing effective breeding schemes. These methods, although providing inspiration for this work, will need to be expanded to be directly applicable to community selection. Thus, it will be important to develop new theories that incorporate unique features of community selection, such as ecological dynamics resulting from species interactions, as well as intra-versus inter-community selection.

## Methods

### Calculating community function landscape *P*(*T*) and the ecological attractor

Our model community is similar to that used in our previous work [15]. The community function landscape plots *P*(*T*) as a function of *ϕ_M_* (0) and 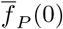. Assume that all cells have the same phenotype (all M cells have the same 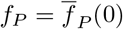), that death and birth processes are deterministic, and that there is no mutation.*P*(*T*) can then be numerically integrated from the following set of scaled differential equations for any given pair of *ϕ_M_*(0) and 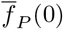 [15]:

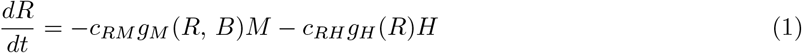

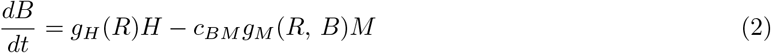

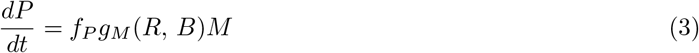

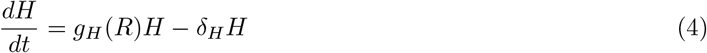

where

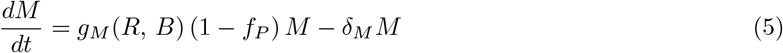

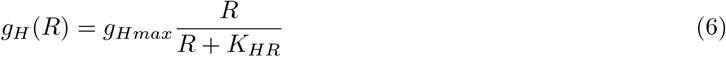

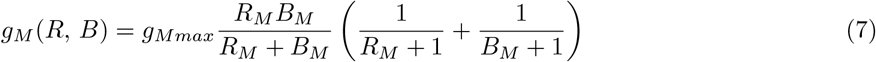

and *R_M_* = *R/K_MR_* and *B_M_* = *B/K_MB_*.

Eq. 1 states that Resource *R* is depleted by biomass growth of M and H, where *c_RM_* and *c_RH_* represent the amount of *R* consumed per unit of M and H biomass, respectively. Eq. 2 states that Byproduct B is released as H grows, and is decreased by biomass growth of M due to consumption (*c_BM_* amount of B per unit of M biomass). Eq. 3 states that Product P is produced as *f_P_* fraction of potential M growth. Eq.4 states that H biomass increases at a rate dependent on Resource *R* in a Monod fashion (Eq. 6) and decreases at the death rate *δ_H_*. Eq. 5 states that M biomass increases at a rate dependent on Resource *R* and Byproduct *B* according to the Mankad and Bungay model (Eq. 7, [43]) discounted by (1 − *f_P_*) due to the fitness cost of making Product, and decreases at the death rate *δ_M_*. In the Monod growth model (Eq. 6), *g_H_max__* is the maximal growth rate of H and *K_HR_* is the *R* at which *g_H_max__*/2 is achieved. In the Mankad and Bungay model (Eq. 7), *K_MR_* is the *R* at which *g_M_max__*/2 is achieved when *B* is in excess; *K_MB_* is the *B* at which *g_M_max__*/2 is achieved when *R* is in excess.

To construct the landscape in Figure 2C, we calculated *P*(*T*) for every grid point on a 2D quadrilateral mesh of 10^−2^ ≤ *ϕ_M_*(0) ≤ 0.99 and 10^−2^ ≤ 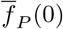 ≤ 0.99 with a mesh size of Δ*ϕ_M_*(0) = 10^−2^ and 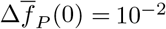. To construct the landscapes in Figure 5B(ii) and B(iii), *P*(*T*) is similarly calculated on a 2D grid with a finer mesh of Δ*ϕ_M_* (0) =5 × 10^−3^ and 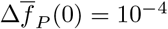.

To calculate the species composition attractor with no spiking, we integrate Eqs. 1–5 to obtain *ϕ_M_*(*T*) − *ϕ_M_* (0) for each grid point on the 2D mesh of *ϕ_M_* (0) and 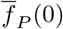. The contour of *ϕ_M_*(*T*) − *ϕ_M_* (0) = 0 is then the attractor under no spiking (orange curve in Figure 3B and Figure 5). Under *x%-H* spiking, *x%* of the biomass in Newborns are replaced with H cells, thus the attractor is the contour of (1 − *x%*)*ϕ_M_*(*T*) − *ϕ_M_*(0) = 0 (teal curve in Figure5). Compared with the orange attractor under no spiking, the teal attractor is shifted down.

### Simulation code of community selection

Simulation codes used in this work are largely similar to those in our previous work, except for the modification to simulate species spiking. Below we briefly recapture the flow of the simulation, which can be found in Ref. [15].

Each simulation begins with *n_tot_* = 100 identical Newborns with 60 M cells and 40 H cells of biomass 1 so that the total biomass of each Newborn is *BM_target_* = 100. Each Newborn is supplied with abundant agriculture waste and a fixed amount of Resource that supports the growth of 10^4^ total biomass. Maturation time is set to be short (*T* = 17, ~6 generations) to avoid Resource depletion (i.e. stationary phase in experiments) and cheater takeover.

Each maturation cycle is divided into 340 time steps of length Δ*τ* = 0.05. During each time step, the biomass of each M and H cell grows according to Eq. 4 and 5 while the concentration of Resource, Byproduct and Product change according to Eq. 1-3. At the end of each Δ*τ*, if a cell’s biomass exceeds the threshold of 2, the cell divides into two identical daughter cells. Each daughter cell then mutates with a probability of *P_mut_*-For a M cell, its *f_P_*, *g_M_max__*, *K_MR_* and *K_MB_* mutate independently. For a H cell, its *g_H_max__* and *K_HR_* mutate independently. In most simulations, only *f_p_* of M mutates while other phenotypes are held at their bounds whose values are shown in the “Bounds” column of Table 1. If a mutation occurs, it could be a null mutation with a probability of 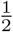. A null mutation reduces *f_P_*, *g_M_max__* and *g_H_max__* to zero, while increases *K_MR_, K_MB_* and *K_HR_* to infinity (equivalent to reducing affinities to zero). If a mutation is not null, it modifies each phenotype by ~5%-6% on average. Specifically, each phenotype is multiplied by (1 + Δ*s*), where Δ*s* is a random variable with a distribution

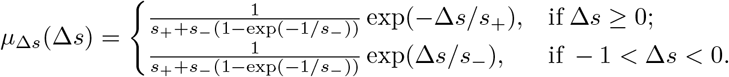

Here, *s*_+_ = 0.05 and *s_−_* = 0.067 are the average percentage by which a mutation increases or decreases a phenotype, respectively (for parameter justifications, see [15]).

At the end of each time step Δ*τ*, each M and H cell dies with a probability of *δ_M_*Δ*τ* and *δ_H_*Δ*τ*, respectively. At the end of a maturation cycle, the amount of Product P in the Adult, *P*(*T*), is the community function. In some simulations, measurement noise is added to the true *P*(*T*) to yield the measured community function. For the simple scenario where only *f_P_* is modified by mutations (e.g., Figure 6(E-H)), measurement noise is a normal random variable with 0 mean and standard deviation of 100, approximately 10% of the ancestral community function. For the complex scenario where 6 phenotypes of H and M are modified by mutations (e.g. Figure 7), measurement noise is a normal random variable with 0 mean and standard deviation of 50, comparable to the ancestral community function. Top *n_chosen_* Adult communities with the highest measured function are chosen to be reproduced. Sometimes more than *n_chosen_* Adults might be needed to obtain *n_tot_* = 100 Newborns for the next cycle if there is not enough biomass in *n_chosen_* Adults.

Chosen Adults are reproduced into Newborns with different methods. If “cell sorting” is used, then Newborn total biomass will be very close to the target *BM_target_* = 100, and species ratio will be very close to that of the parent Adult. If “pipetting” is used, then Newborn total biomass or species composition are allowed to fluctuate stochastically. If fraction *φ_s_* of a Newborn’s biomass is to be replaced by M or H cells, each Newborn gets on average a biomass of *BM_target_* (1 − *φ_s_*) from its parent Adult community and on average a biomass of *BM_target_φ_S_* from M or H spiking mix. Specifically, suppose that the biomass of an Adult is *BM* (*T*) = *M*(*T*)+ *H*(*T*) where *M*(*T*) and *H*(*T*) are the biomass of M and H at time *T*, respectively. If a fraction *φ_s_* of each Newborn’s biomass is to be replaced by a spiking mix, this Adult is then reproduced into *n_D_* Newborns, where *n_D_* = ⌊*BM*(*T*)/[*BM_target_* (1 − *φ_s_*)]⌋ and ⌊*x*⌋ is the floor (round down) function. If *n_D_* is larger than *n_tot_*/*n_chosen_*, only *n_tot_*/*n_chosen_* Newborns are kept. Otherwise, all *n_D_* Newborns are kept and as many additional Adults with the next highest functions are reproduced to obtain *n_tot_* Newborns for the next cycle. These Newborns are then topped off with either M or H spiking mixes so that their total biomass are on average *BM_target_* = 100. Note that the fold of dilution of an Adult is calculated based on biomass, a continuous variable. However, the biomass is composed of individual biomass of discrete cells. During reproduction, integer number of cells are distributed into each Newborn communities.

### Community reproduction with species spiking in the simple scenario when only *f_P_* is modified by mutations

In the simple scenario where only *f_P_* of M is modified by mutations, phenotypes of all H cells are the same. Within an Adult community, all H cells also have identical individual biomass *L_H_*, because simulations start with H cells of biomass 1 and because growth is synchronous. To mimic reproducing a chosen Adult through sorting a fixed biomass into each Newborn with a *φ_S_*-H spiking strategy, H and M cells from the chosen Adult are randomly assigned to a Newborn community until its total biomass came closest to *BM_target_* (1 − *φ_s_*). If *φ_S_ >* 0, the number of H cells supplemented to the Newborn community is the nearest integer to 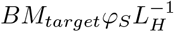. Because integer number of cells are assigned to each Newborn, the total biomass might not be exactly *BM_target_* but with a small deviation of ~2 biomass unit.

To mimic reproducing through pipetting, each M and H cell in an Adult community is assigned a random integer between 1 and dilution factor *n_D_*. All cells assigned with the same random integer are then dealt to the same Newborn, generating *n_D_* Newborn communities. To replace *φ_s_* of a Newborn with H biomass, the number of H cells distributed into each Newborn is a random number drawn from a Poisson distribution of a mean of 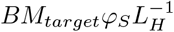.

### Community reproduction in the complex scenario when both H and M cells evolve

In the more complex scenario, both H and M evolve. We thus need to spike with evolved H and M clones. Additionally, Newborns are spiked with H or M clones from their own lineage as demonstrated in Figure S4A. Below we describe the the simulation code for the experimental procedure (Figure S4A) we simulated.

Specifically, after H and M cells from a chosen Adult in Cycle *C −* 1 are randomly assigned to generate Newborns, a pre-set number of H or M cells are randomly picked from the remaining of this Adult. At the end of Cycle *C*, we choose 10 Adults with highest functions. Each Adult is then reproduced through pipetting with *φ_S_*-H spiking strategy. Since each chosen Adult usually gives rise to 10 Newborns, the number of cells distributed from the chosen Adult to each Newborn is drawn from a multinomial distribution. Specifically, denote the integer random numbers of cells that would be assigned to 10 Newborns to be {*x*_1_,*x*_2_, … *x*_10_}. If the chosen Adult has a total biomass of *BM*(*T*), and *I_M_* M cells and *I_H_* H cells (both *I_M_* and *I_H_* are integers), the probability that {*x*_1_, *x*_2_, … *x*_10_} cells are assigned to 10 Newborns respectively, and *x*_11_ cells remain, is

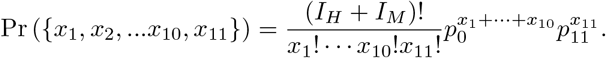

Here, *p_0_* = *BM_target_* (1 − *φ_s_*) /*BM*(*T*) is the probability that a cell is assigned to one of 10 Newborns, *p_11_* = 1 − 10*p*_0_ is the probability that a cell is not assigned to Newborns. Thus 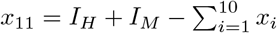 is the number of cells remaining after reproduction, from which H and M cells are randomly picked to generate the spiking mix for the next cycle.

These 10 Newborns are then spiked with either H or M cells. Suppose that the current spiking strategy is *φ_S_*-H, then the H biomass spiked into each Newborn is on average *BM_targetφ_S__* so that the total biomass of Newborns is on average *BM_target_*. Suppose that 5 H cells from the parent Adult’s lineage are randomly picked at the end of Cycle C-1, and that they have biomass {*L*_*H*_1__, *L*_*H*_2__, *L*_*H*_3__, *L*_*H*_4__, *L*_*H*_5__}, respectively. The total number of H cells assigned to each Newborn, *x_H_*, is then randomly drawn from a Poisson distribution with a mean of 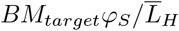, where 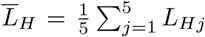 is the average biomass of the 5 H cells. Each spiked H cell has an equal chance of being one of the 5 cells.

### Dynamic spiking strategy

In the complex scenario where 6 phenotype mutate, we estimate heritability of community function under different spiking strategies through parent-offspring regression. Heritability evaluation is carried out about every 100 cycles. Based on the heritability under all candidate spiking strategies, the current spiking strategy is updated if an alternative spiking strategy confers significantly higher community function heritability. At early selection cycles, heritability evaluation can be postponed until within-community selection improve cell growth sufficiently to provide sufficient biomass for heritability evaluation.

During one round of heritability evaluation, heritability of community function is estimated through parent-offspring community function regression under all candidate spiking strategies (Figure S4B). To evaluate heritability under one spiking strategy, 100 Newborn communities are generated under this spiking strategy. After these mature into Adults, their functions are the parent functions. Each Adult parent then gives rise to 6 Newborn offspring under the same spiking strategy. When the 6 Newborn offspring mature into Adults, the median of their functions is the offspring function.

When offspring functions are plotted against their parent functions, the slope of the least squares linear regression (green dashed line in Figure S4B) is the heritability of community function. Heritability of a community function is thus similar to heritability of an individual trait, except that we use median instead of mean of offspring functions, because median is less sensitive to outliers. The 95% confidence interval of heritability is then estimated by non-parametric bootstrap [44, 45]. More specifically, first, 100 pairs of parent-offspring community functions are re-sampled with replacement. Second, heritability is calculated with the re-sampled data. Third, 1000 heritabilities are calculated from 1000 independent re-samplings, from which the 95% confidence interval is estimated from the 5th and 95th percentile. An alternative spiking strategy is considered more advantageous than the current spiking strategy if heritability of the alternative spiking strategy is higher than the right endpoint of the 95% confidence interval of the heritability of the current spiking strategy. If more than 1 alternative spiking strategies are more advantageous, the one with the highest heritability is implemented to replace the current strategy. Similarly, an alternative spiking strategy is considered more disadvantageous if heritability of the alternative spiking strategy is lower than the left endpoint of the 95% confidence interval of the heritability of the current spiking strategy. When implementing random spiking strategy, the current spiking strategy is updated with a strategy randomly picked from candidate spiking strategies.

## Supporting Figures

**Figure S1:**
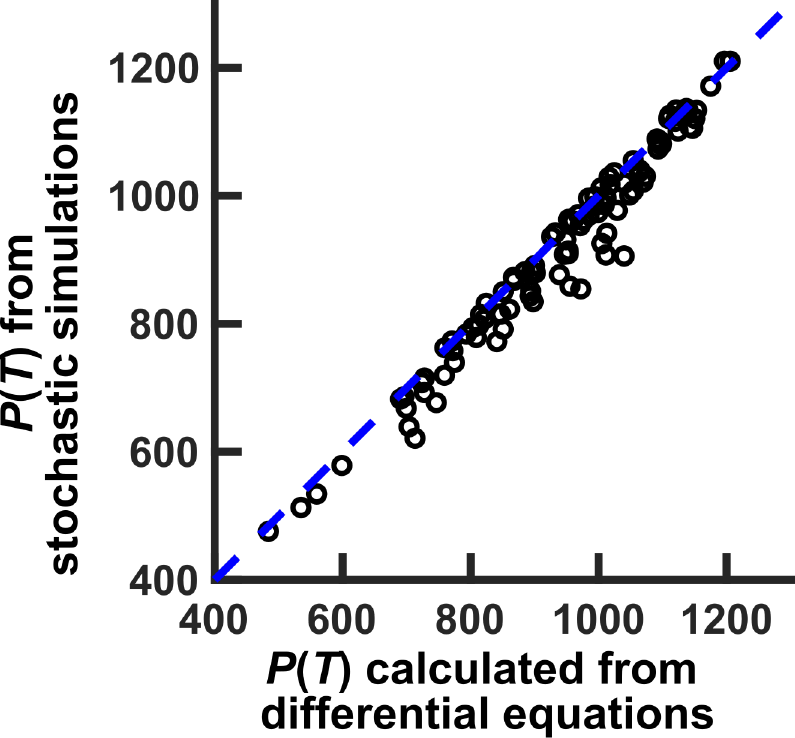
Community functions *P*(*T*) obtained from stochastic simulations can be adequately predicted by numerically integrating deterministic differential Eq. 1-5. We calculated *P*(*T*) of 100 communities from the 2000th cycle of the simulation shown in black curves in Figure 7(A-B) with differential equations, and compared community functions to those obtained from stochastic simulations. For each community, determinants are calculated as the within-Newborn average of M’s cost 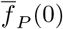, average maximal growth rates 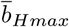 and 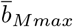, average affinity of M to Byproduct 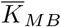, average affinity of M to Resource 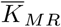, and average affinity of H to the Resource 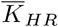.

**Figure S2:**
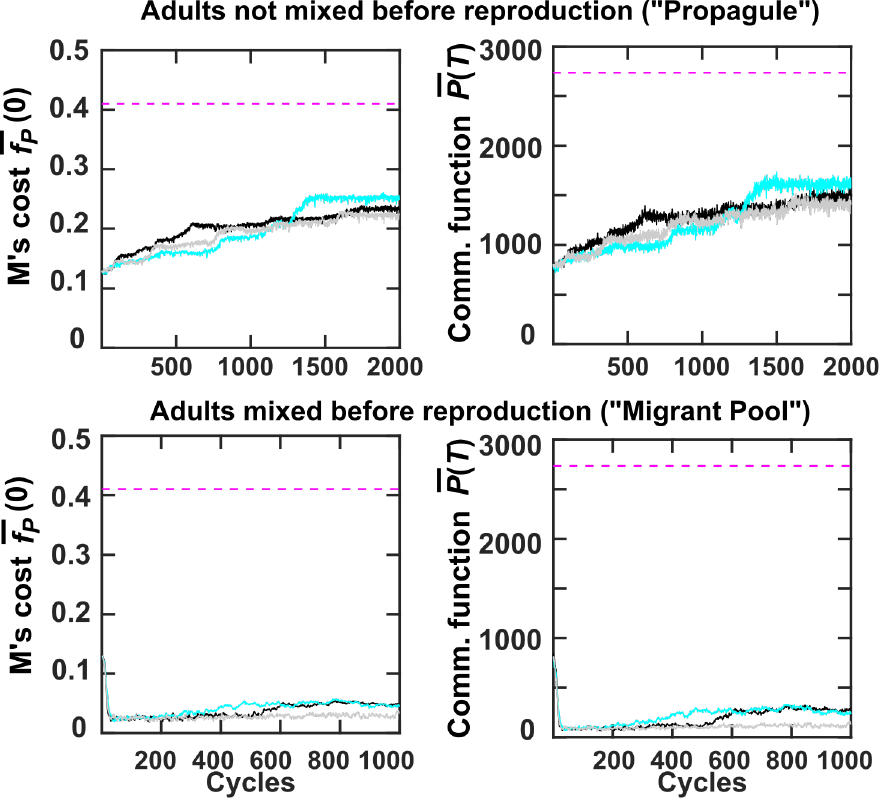
Community reproduction through “propagule” and “migrant pool” methods yield different outcomes. Top 10 communities were chosen for reproduction via pipetting so that Newborn total biomass and species composition stochastically fluctuated. Measurement noise was not considered. Top: Adults were not mixed before reproduction, and each gave birth to 10 Newborns. The data are identical to Fig 3 G and H from [15]. Bottom: Adults were mixed, and the mixture gave birth to 100 Newborns

**Figure S3:**
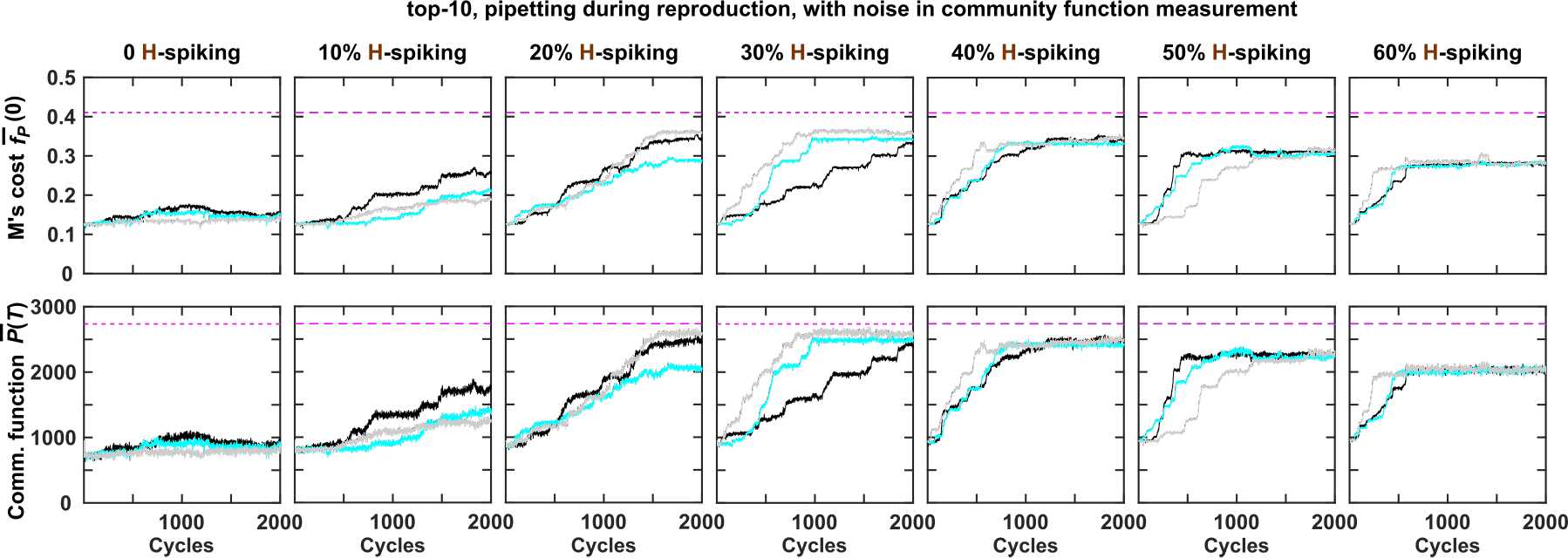
Spiking H at a wide range of percentages improves selection outcome. Adults ranking top 10% in measured community functions (= true community function + a normal random variable with zero mean and standard deviation of 100 ~10% of the ancestral community function) were chosen to reproduce. Each of the 10 chosen Adults reproduced 10 Newborns. The total biomass of a Newborn fluctuated around the target value (*BM_target_* = 100), of which the indicated percentage was “pipetted” from an H monoculture while the rest was “pipetted” from the parent Adult. Consequently, Newborn total biomass and species composition fluctuated stochastically according to sampling error. Top panels show the dynamics of heritable determinant 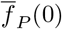, the cost paid by M averaged first within and then among chosen Adults. Bottom panels show 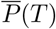, the average community function of chosen Adults. Black, cyan and gray curves represent three independent replicates.

**Figure S4:**
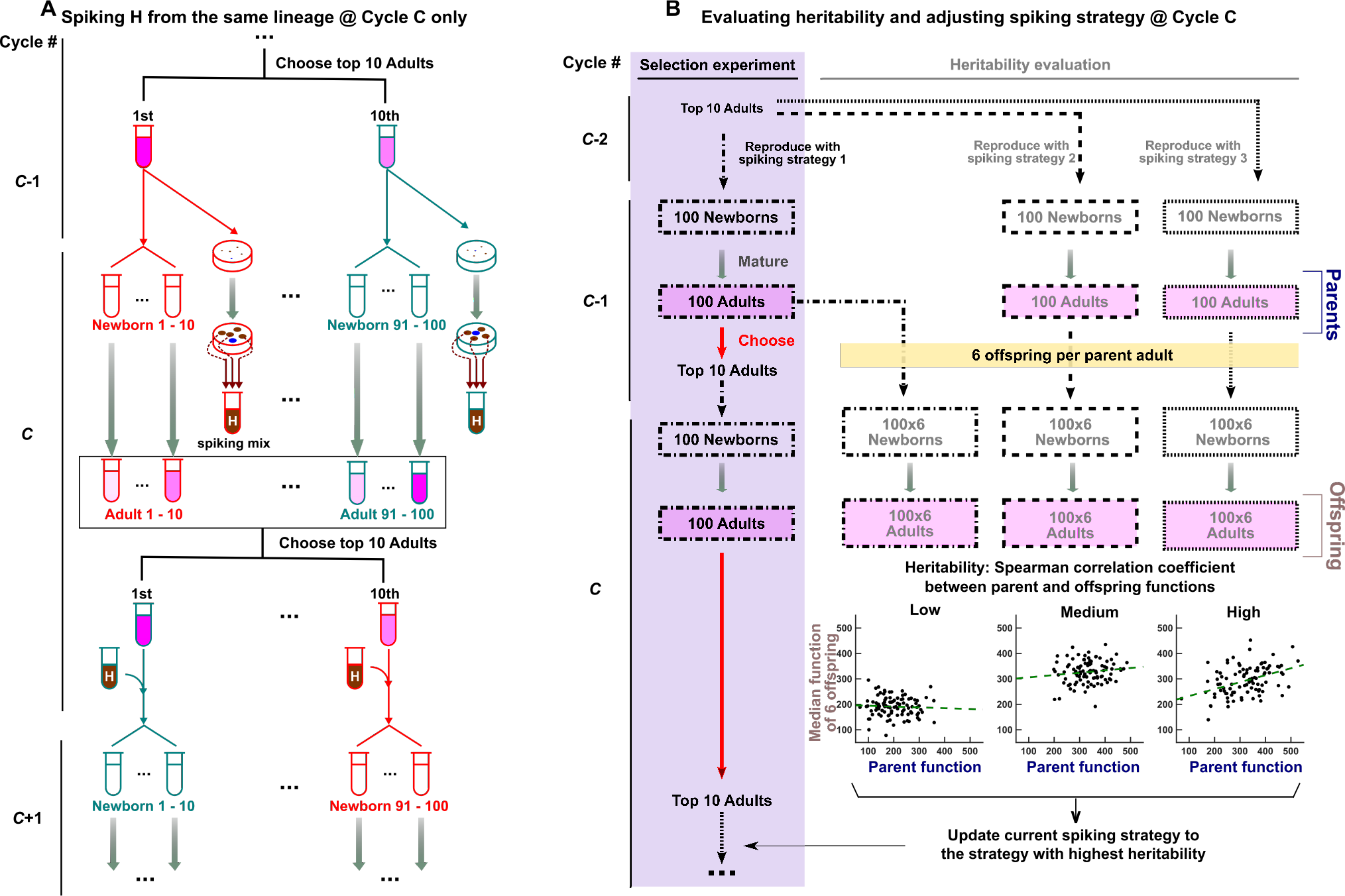
Schematics of dynamic spiking. **(A)** Spiking during community reproduction at Cycle *C* with H colonies isolated at Cycle *C* − 1. Cycle *C* − 1: Each chosen Adult in Cycle *C* − 1 is outlined in different colors to indicate separate lineages. After each chosen Adult gives rise to 10 Newborns, the remaining Adult is plated (in simulations, this means that 5 cells are randomly chosen from the remaining Adult). Communities and the plates share identical color outline to indicate their shared lineage. Cycle *C*: The 100 Newborns mature into Adults and H colonies develop on the plates. Several H colonies are randomly chosen and mixed at equal biomass as the “spiking mix”. At the end of Cycle *C*, all Adults are ranked according to their function. To reproduce a chosen Adult, offspring Newborns are spiked with the spiking mix from the same lineage (same outline color). **(B)** Evaluating heritability and adjusting spiking strategy at Cycle *C*. Here we illustrate 3 candidate spiking percentages, with the purple box highlighting the current spiking strategy. Cycle *C* − 2: The 10 chosen Adults are reproduced with all 3 spiking strategies, generating 3 sets of 100 Newborns (different line styles). Cycle *C* − 1: The 3 sets of 100 Newborns mature into Adults, and their functions define the “Parent function” (blue axis label of the scatter plot). Each Adult then gave birth to 6 offspring Newborns under the respective spiking strategy, and when these mature, the median of 6 functions defines the “Offspring function” (brown axis label of the scatter plot). Cycle *C*: At the end of cycle *C*, we can scatter plot the function of each parent against the average function of its 6 offspring. The heritability is estimated from the slope of the least squares regression line. In parallel, the current spiking strategy continues (purple box) until Cycle *C*, where spiking strategy is updated. In this example, the updated current strategy is spiking strategy 3.

**Figure S5:**
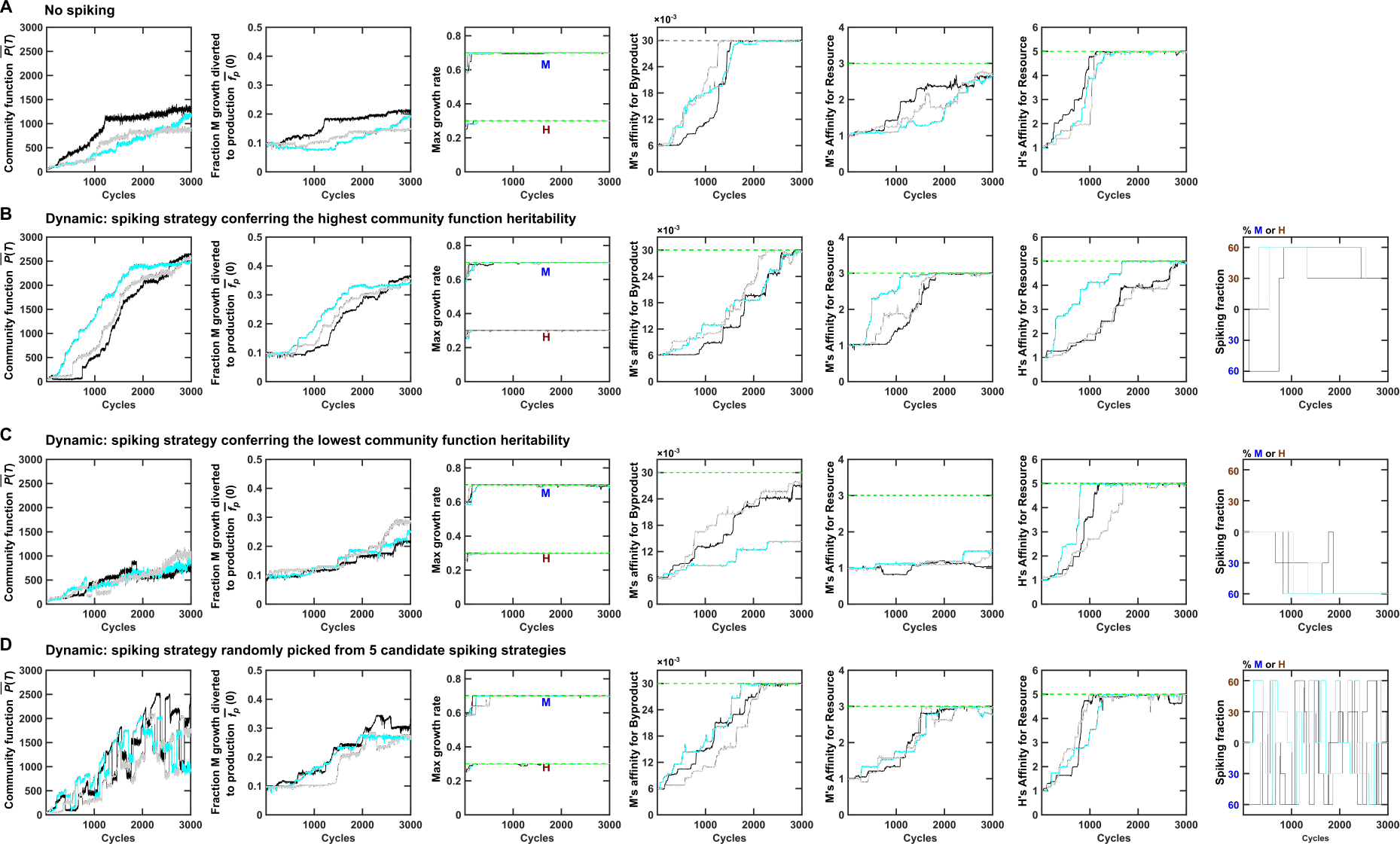
Evolution dynamics of artificial community selection under no-spiking strategy **(A)** and different dynamic spiking strategies **(B-D)**. Spiking strategies conferring the highest community function heritability result in the best outcome (Compare panel B vs. panels A, C and D). The first two columns of A and B are the same as Figure 7. Spiking mix used in (B-D) are expanded from 5 clones of M or H. Black, cyan and gray curves represent 3 independent simulation replicas. The first column displays the dynamics of the community functions averaged among the 10 chosen Adults. Columns 2-6 display the dynamics of the following community function determinants averaged among the chosen communities: Newborn average cost paid by M 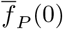, Newborn average maximal growth rates of H and M 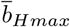 and 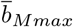, Newborn average affinity of M to Byproduct 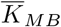, Newborn average affinity of M to Resource 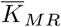, and Newborn average affinity of H to Resource, 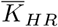. The overbar represents averaging first within each community, then across chosen communities. The last column of (B-D) displays the spiking percentage implemented in corresponding selection simulation.

**Figure S6:**
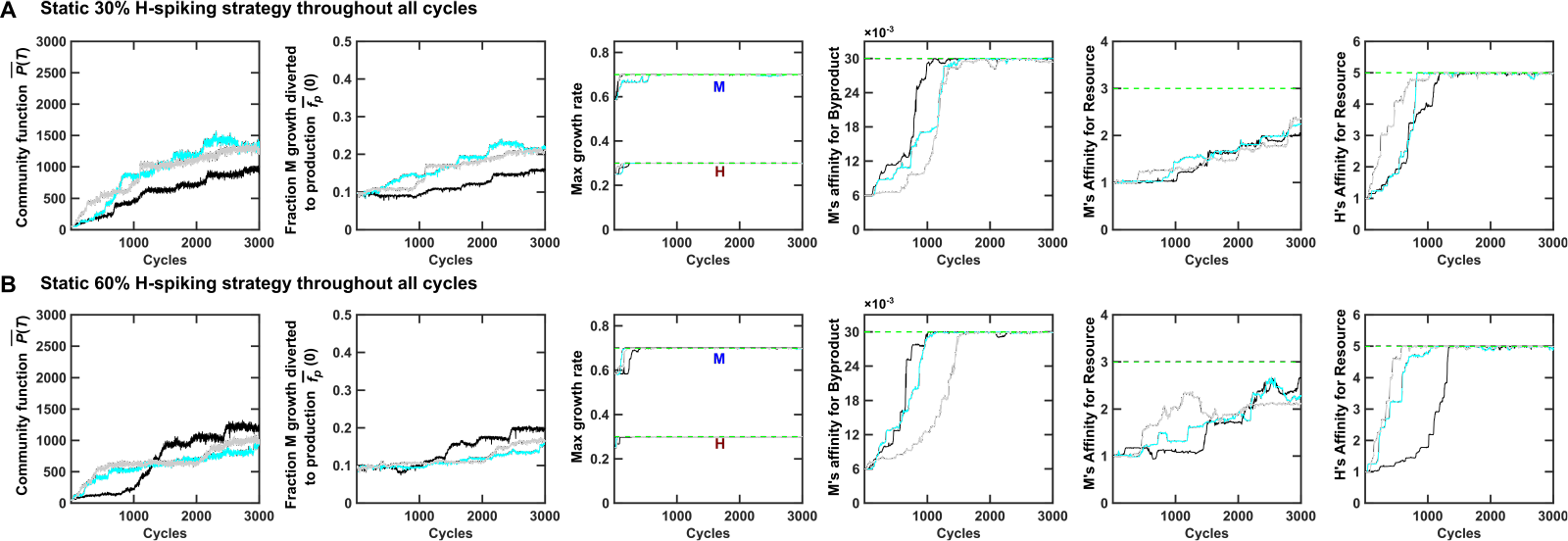
Static spiking strategy does not work as well as dynamic spiking. Compared to dynamic spiking (Figure 7B), static spiking was not as effective. **(A)** static 30%-H spiking strategy. **(B)** static 60%-H spiking.

**Figure S7:**
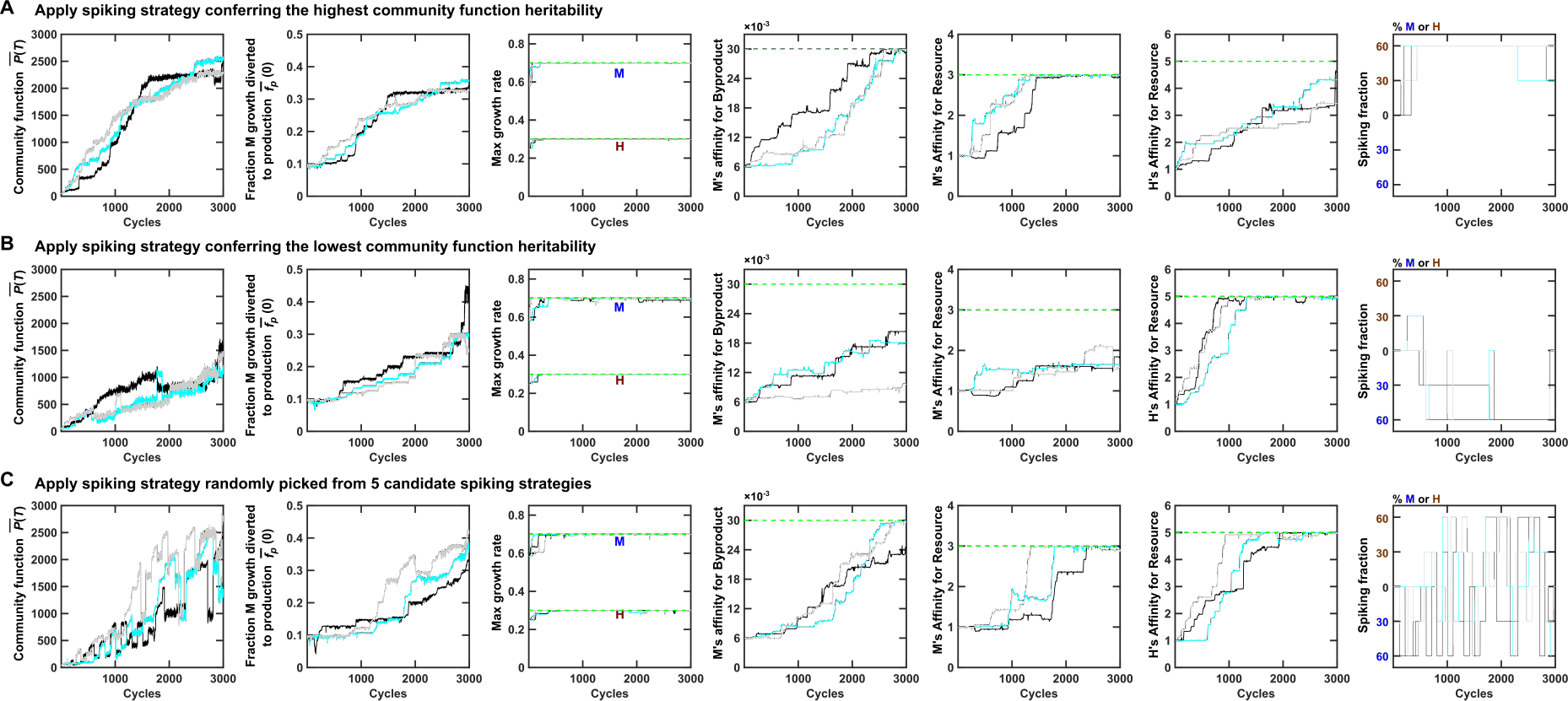
Spiking with 1 clone of H or M generated qualitatively similar dynamics as spiking with 5 clones of H or M (Figure S5). Figure legends are the same as Figure S5.

**Figure S8:**
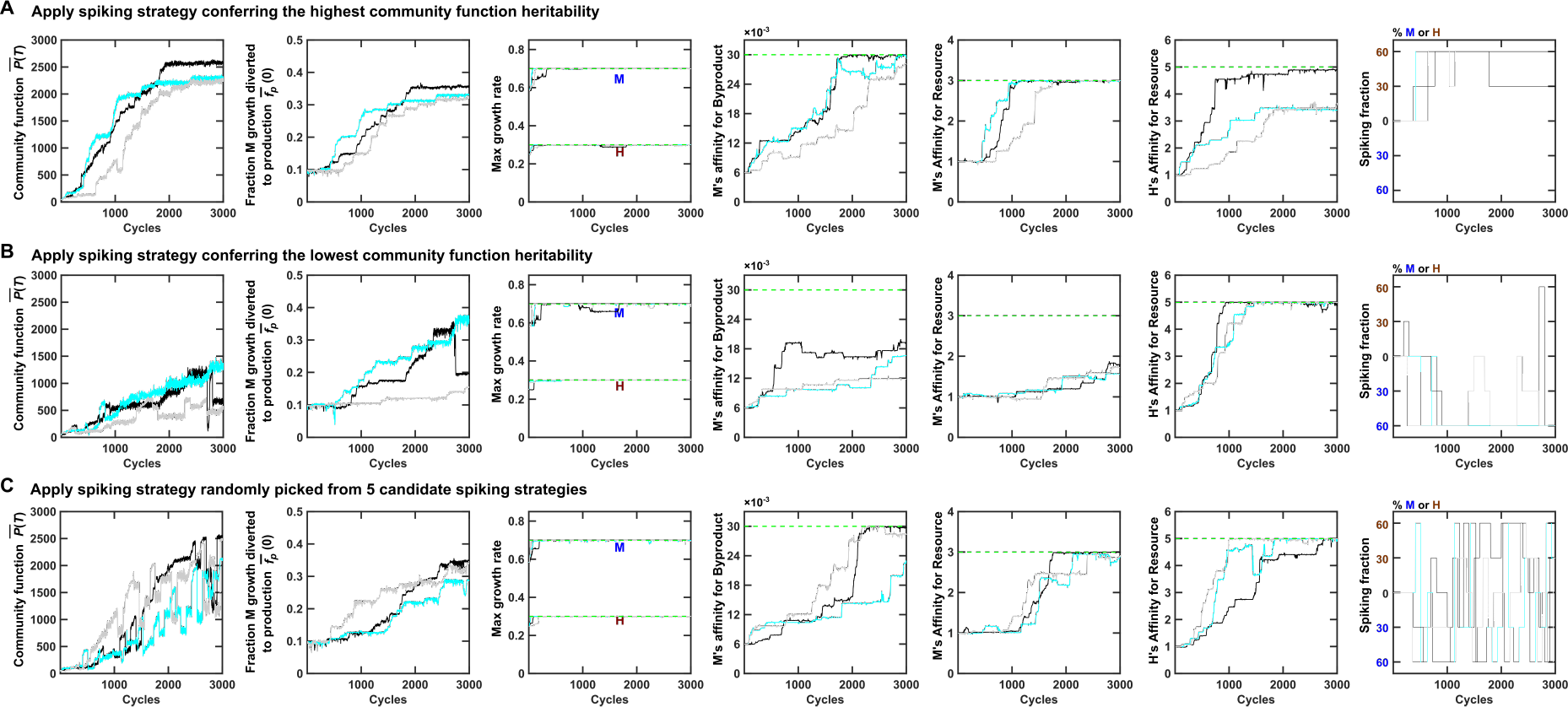
Spiking with 2 clones of H or M generated qualitatively similar dynamics as spiking with 5 clones of H or M (Figure S5). Figure legends are the same as Figure S5.

**Figure S9:**
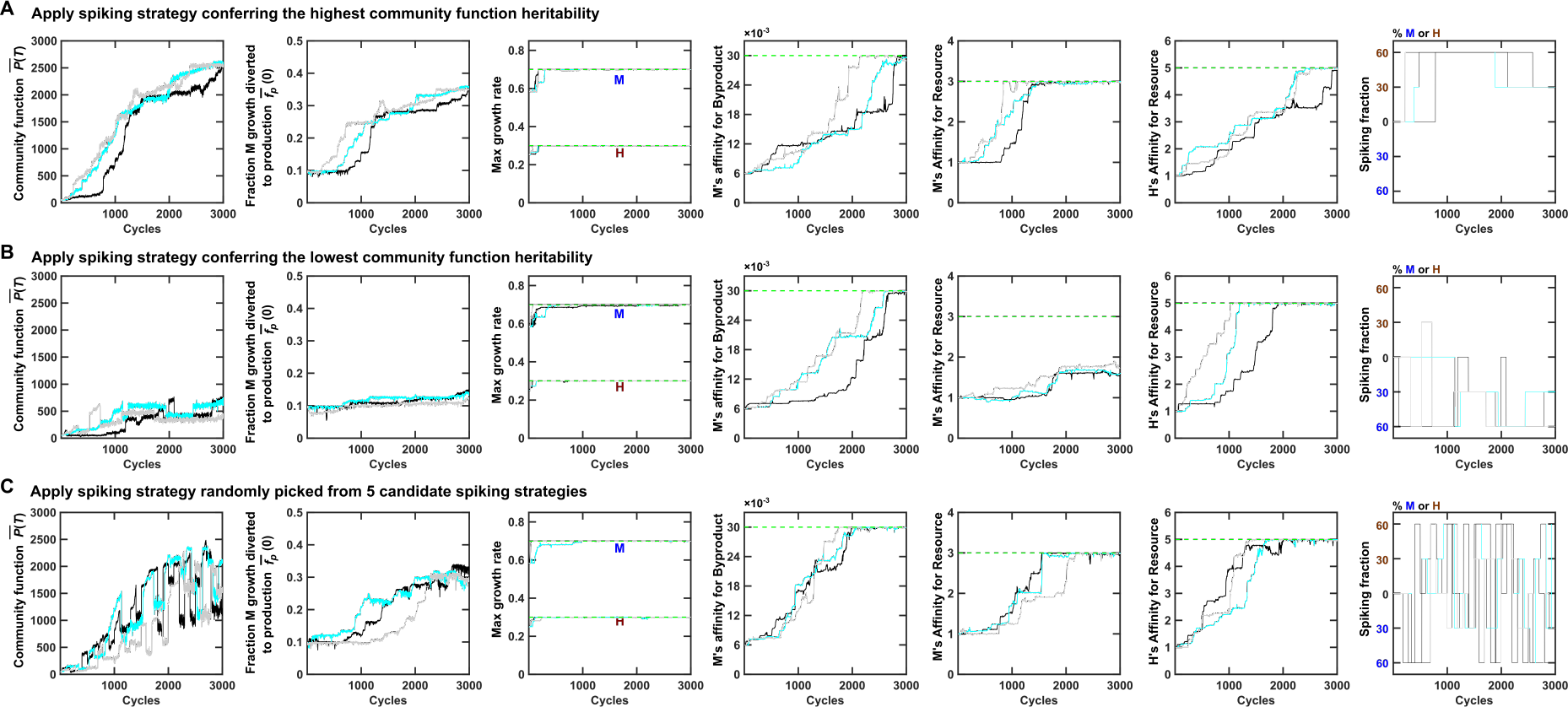
Spiking with 10 clones of H or M generated qualitatively similar dynamics as spiking with 5 clones of H or M (Figure S5). Figure legends are the same as Figure S5.

**Figure S10:**
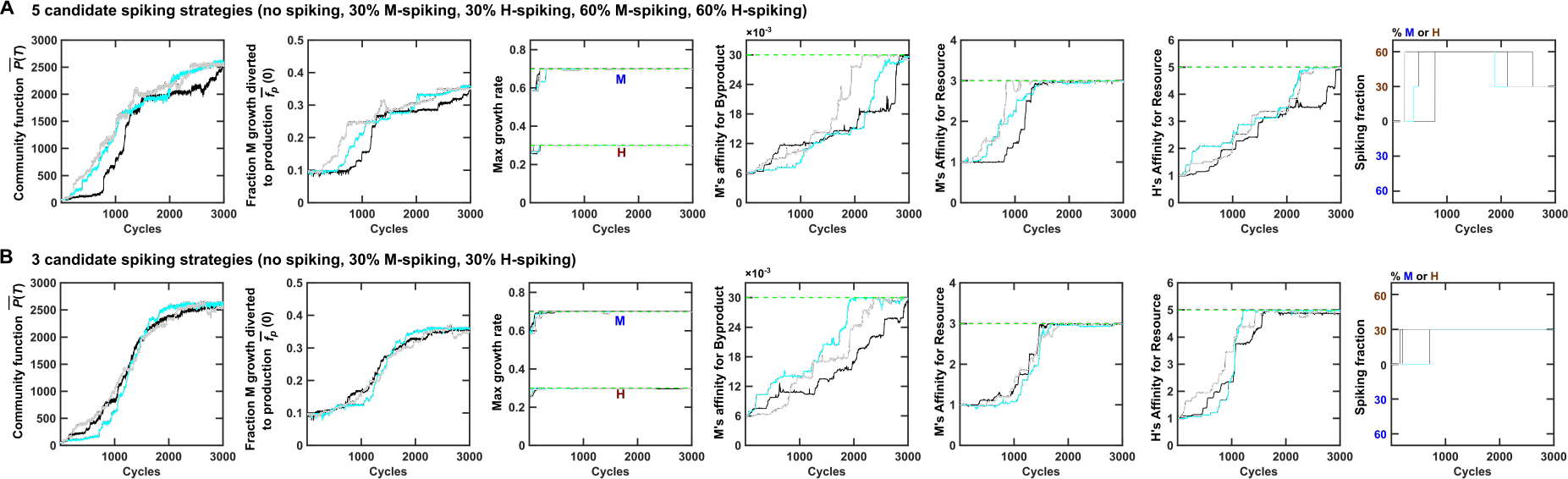
Evolution dynamics of community selection under the dynamic spiking strategy with 5 or 3 candidate spiking strategies. Spiking mix consisted of 10 evolved H clones or 10 evolved M clones. The legends are the same as Figure S5.

**Figure S11:**
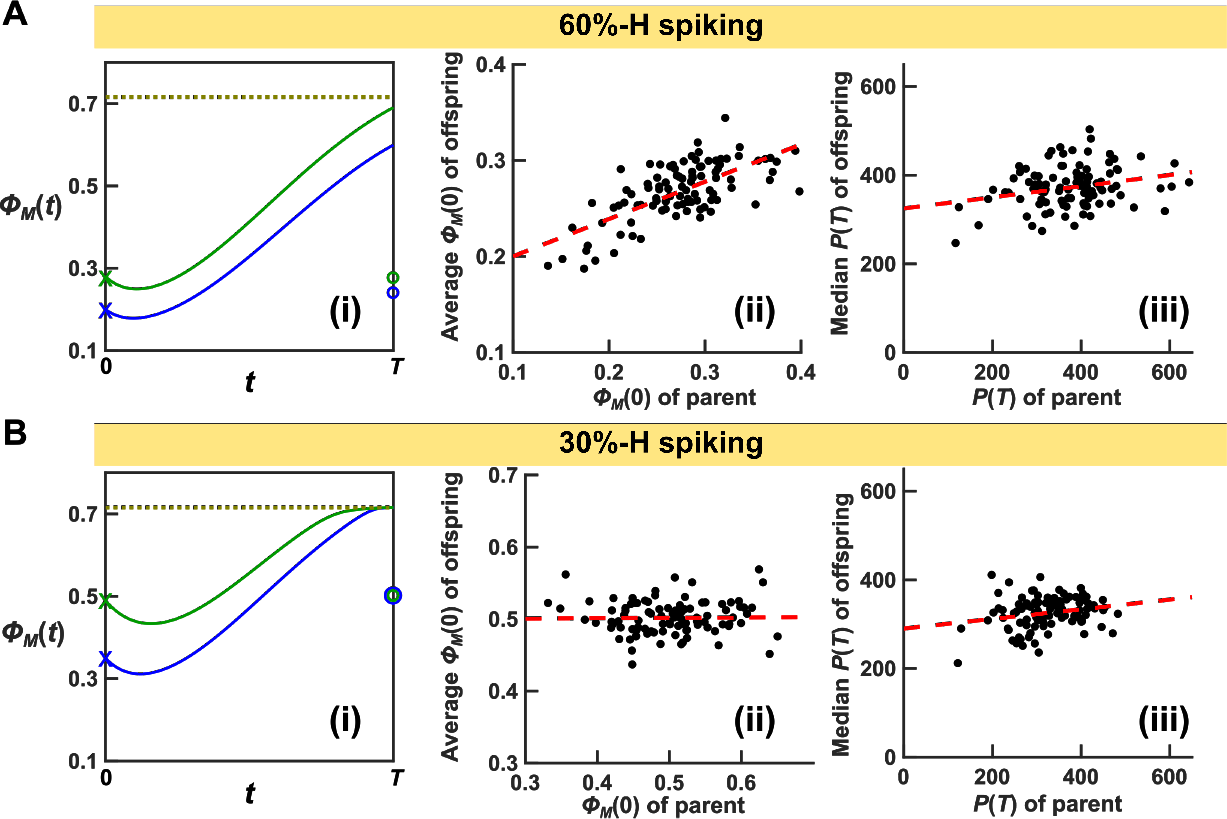
Spiking a species at too high a fraction can make a non-heritable determinant seem heritable. Compare 60%-H spiking **(A)** versus 30%-H spiking **(B)**. **(i)** Species composition dynamics of two communities (blue and green). Starting at Newborns immediately after spiking (crosses; parent Newborns), species compositions moves toward the attractor (dotted line). In 30%-but not 60%-H spiking, species composition reaches the attractor by the end of maturation. After reproducing with respective spiking percentages, average offspring Newborn compositions are plotted with circles. In 30%-H spiking, average offspring Newborns have the same species composition (green circle overlapping with blue circle). In contrast, in 60%-H spiking, average offspring Newborns have different species composition (green circle above blue circle). **(ii)** In 60-but not 30%-H spiking, the determinant *ϕ_M_* (0) displays heritability. **(iii)** Consequently, although 60%-H spiking strategy confers similar community function heritability as 30%-H spiking strategy (similar slopes of the red dashed lines), community function improved faster under 30%-H spiking strategy (Figure S10B) than under 60%-H spiking strategy (Figure S10A). This is because in 60%-H spiking, a portion of community function heritability originates from the misleading heritability created by a nonheritable determinant.

